# Invasive cancer and spontaneous regression two weeks after papillomavirus infection

**DOI:** 10.1101/2024.09.04.611275

**Authors:** Andrea Bilger, Ella T. Ward-Shaw, Denis L. Lee, Renee E. King, Michael A. Newton, Darya Buehler, Kristina A. Matkowskyj, John P. Sundberg, Rong Hu, Paul F. Lambert

## Abstract

Development of invasive cancer in mammals is thought to require months or years after initial events such as mutation or viral infection. Rarely, invasive cancers regress spontaneously. We show that cancers can develop and regress on a timescale of weeks, not months or years. Invasive squamous cell carcinomas developed in normal adult, immune-competent mice as soon as 2 weeks after infection with mouse papillomavirus MmuPV1. Tumor development, regression or persistence was tissue- and strain-dependent. Cancers in infected mice developed rapidly at sites also prone to papillomavirus-induced tumors and cancers in humans – the throat, anus, and skin – and their frequency was increased in mice constitutively expressing the papillomavirus E5 oncogene, which MmuPV1 lacks. Cancers and dysplasia in the throat and anus regressed completely within 4-8 weeks of infection; however, skin lesions in the ear persisted. T-cell depletion in the mouse showed that regression of throat and anal tumors requires T cells. We conclude that papillomavirus infection suffices for rapid onset of invasive cancer, and persistence of lesions depends on factors including tissue type and host immunity. The speed of these events should promote rapid progress in the study of viral cancer development, persistence, and regression.

**Summary Graphic:** 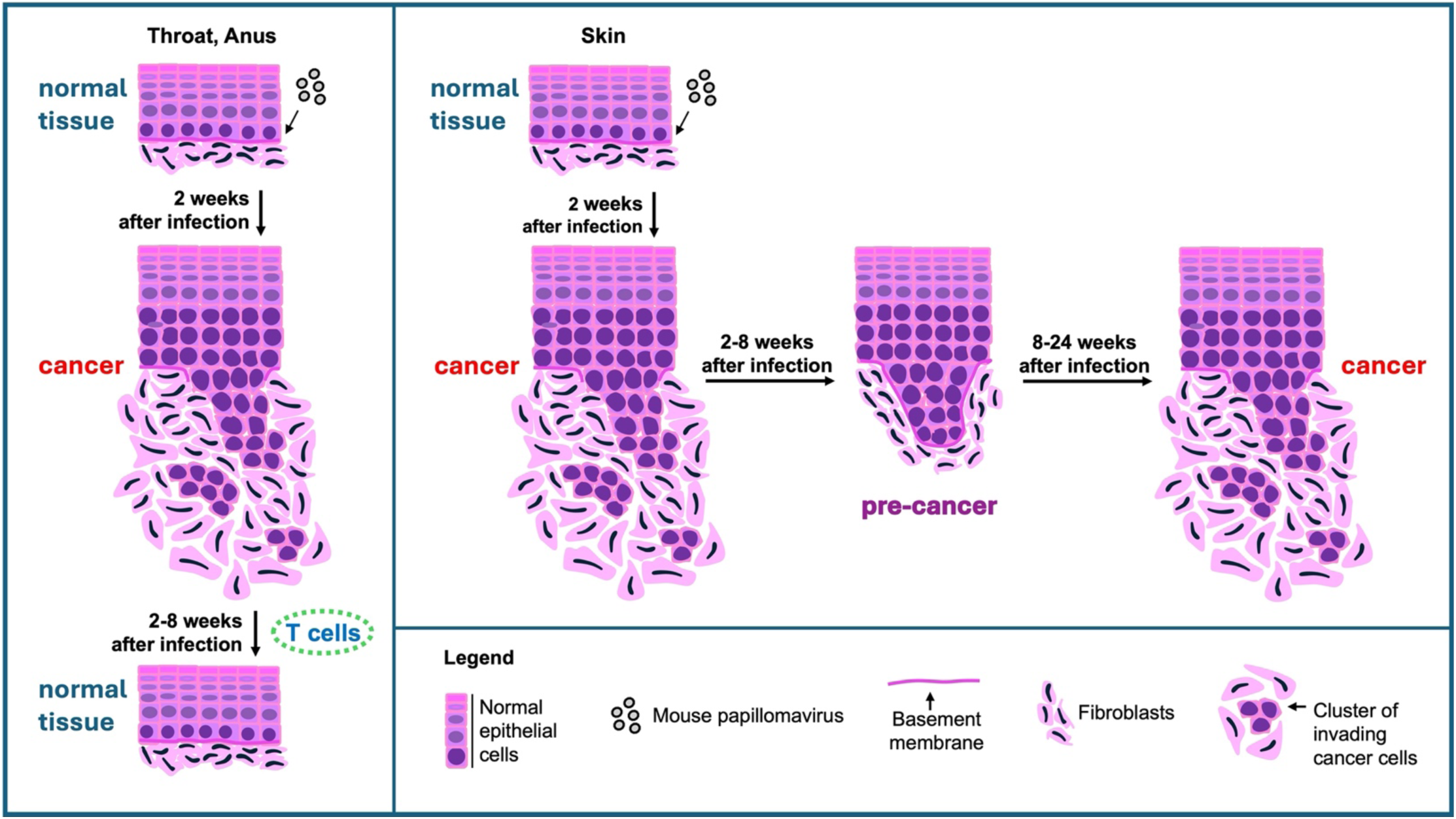

---

Assessing the speed at which cancer develops in adult humans from a cell or cells that incur an initiating insult is difficult because the time of the initial damage is often not known. However, viral infection, responsible for approximately 12% of cancers worldwide^1^, can be monitored. Infections rarely lead to cancer, and years frequently pass between infection and detection of invasive cancer. Human Papillomaviruses (HPVs) are responsible for approximately 5% of cancers worldwide^1^. The median age of HPV infection that leads to cancer is estimated to be 21 years; the median age of cancer detection, depending on the tissue, is 50-68 years^2,3^. This lag between infection and detection of cancer suggests that cancer generally develops slowly from infected cells^4^.

Harald zur Hausen, citing this time lag, stated that “no human cancer arises as the acute consequence of infection,” a view currently still favored^5,1^. However, some observations suggest infection alone can cause cancer to arise rapidly. The development of lymphoma and other abnormal lymphoproliferation frequently occurs in chemically immune-suppressed organ transplant recipients who are seronegative for Epstein Barr Virus (EBV) prior to the transplant operation. These patients develop primary EBV infections post transplantation, with EBV likely sourced from the donor’s tissue^6,7,8^. Cases of lymphoma in these patients have been detected as soon as 2 months post transplantation^6^. Human cord blood cells infected with EBV and injected immediately into the peritoneum of immune-deficient mice yield post-transplant-like lymphomas in as little as 4 weeks^9^. The risk for HPV-associated cancers, like those caused by EBV, also increases dramatically with immune suppression, leading to the recommendation that organ transplant recipients be screened for cervical cancer every 6 months in the first year post transplantation^10^. Here, however, it is difficult to distinguish whether fast-developing cervical cancers arise from pre-existing, persistent infections or new infections.

Clinical findings among immune-competent patients also hint that invasive cancers do not always derive from slowly evolving, enlarging benign tumors. Primary cancers associated with head and neck lymph node metastases are often so small they are hard to find. Robotic surgery directed at the base of the tongue (BoT) has shown that many undetected primary cancers are small, invasive HPV-positive cancers in the lingual tonsils^11^. Metastatic cancers of only 2 and 3 mm at the BoT have been detected^12,13^. Notably, one of these small oropharyngeal cancers regressed completely, spontaneously, following biopsy and tonsillectomy^13^. No mechanism has been established for spontaneous cancer regression, which is very rare^14,15^. Robust animal models of the regression of cancers that develop *in situ* (autochthonous cancers, as opposed to cancers that develop from grafts) are limited to swine strains that develop melanoma congenitally or just after birth^16^.

Human and other mammalian papillomavirus infections can cause benign papillomas or squamous cell carcinomas and other invasive cancers that are specific to the papillomavirus genotype, anatomic site, and host species^17^. The study of papillomavirus-induced neoplastic disease received a boost from the discovery of papillomavirus MmuPV1, which infects laboratory mice^18,19,20^. MmuPV1 induces benign papillomas as well as malignant tumors within months of infection in many of the tissues where human papillomaviruses cause disease, such as the oropharynx (part of the throat), the reproductive tract, and the skin^19,20^. Recent studies in the female reproductive tract and the larynx have shown that MmuPV1 can induce moderate to severe dysplasia 1 to 2 weeks post infection^21,22^.

## Papillomavirus causes cancer within 2 weeks in wild-type adult mice

HPV infection is highly associated with oropharyngeal squamous cell carcinoma^23^. The oropharynx is the part of the throat that opens onto the oral cavity; it includes the base of the tongue (BoT). To study development of papillomavirus-induced oropharyngeal disease, adult, immune-competent, wild-type FVB (FVB/NTac) mice were infected at the BoT. A Greer Pick was used to injure the epithelium and deliver ∼10^9^ viral genome equivalents (VGE) of mouse papillomavirus MmuPV1^24^ (Prep 1; Fig 1a; Extended Data Fig. 1a), and tissue was collected 2 weeks post infection (w.p.i.). Disease was assessed independently by one or two pathologists blinded to treatment (R.H. and, for a subset of tissues, J.P.S.).

**Figure 1.**
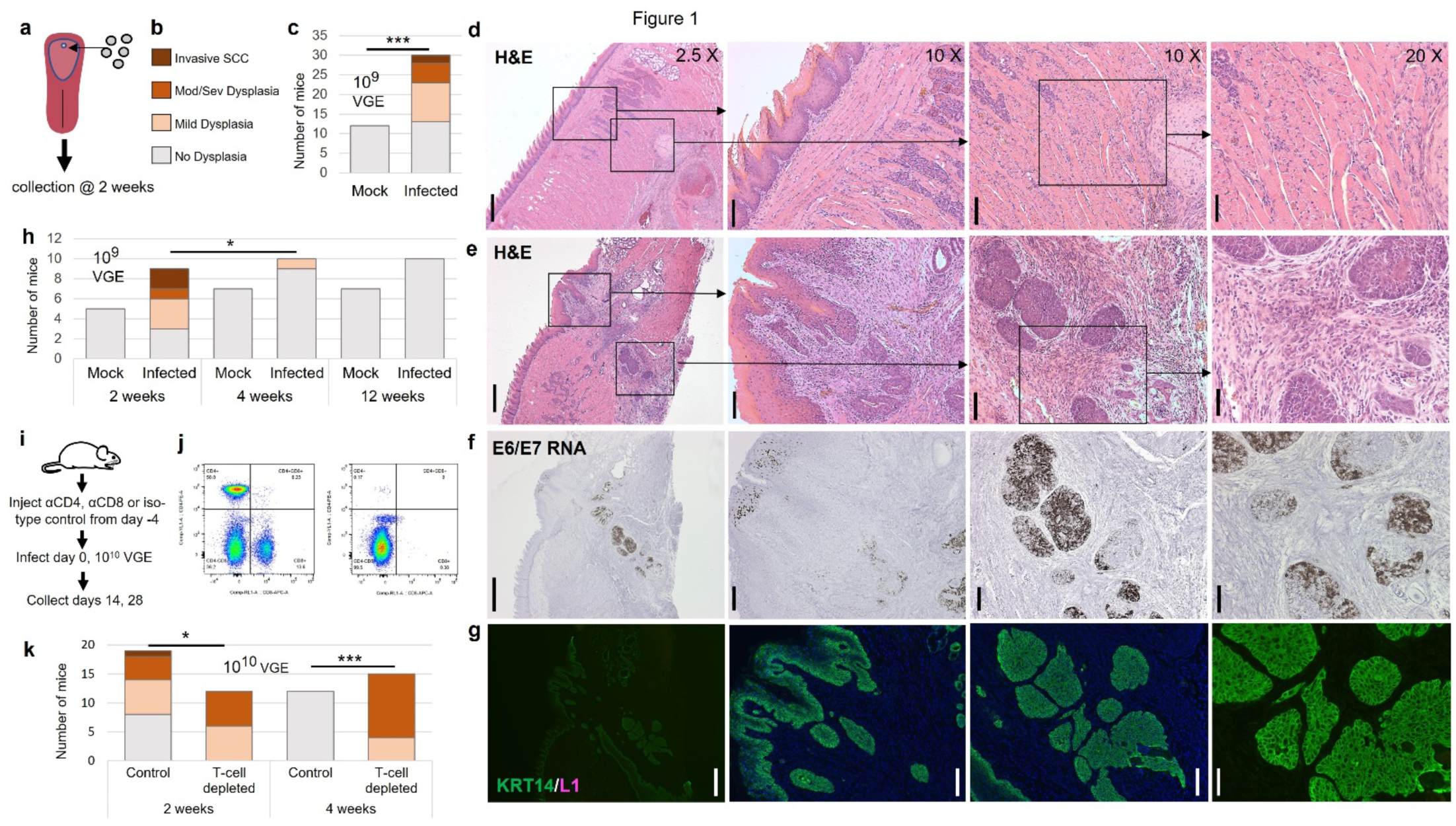
Lesion development and regression in FVB mice. **a,** Illustration showing mouse tongue and location at the BoT to which virus (or PBS) was delivered using the Greer Pick. Tongue tissue was collected 2 w.p.i. **b,** Legend for bar graphs in **c**,**h**,**k**. **c,** Lesion severity 2 weeks post mock infection with PBS (“Mock”) or infection with ∼10^9^ viral genome equivalents (VGE) of MmuPV1 (“Infected”); these data combine the results of 3 non-contemporaneous experiments. Two-sided Wilcoxon Rank Sum (WRS) test of difference in lesion severity: *** p<10^-3^. **d,** BoT mock-infected with PBS, stained with H&E. Panels are images of a single section from one mouse. 10X objective panels: left panel shows surface epithelium and tissue below, including muscle and serous salivary glands; right panel shows adjacent, deeper (ventral) serous salivary gland and muscle tissue. 20X objective panel shows central portion of adjacent 10X panel. Scale bars: 400 um (2.5X objective), 100 um (10X objective), 50 um (20X objective). **e,** Invasive SCC at BoT infected with 10^9^ VGE of MmuPV1, stained with H&E. Panels and scale bars as in **d**, with right two panels focusing on islands of invasive cancer epithelial cells. **f,** Section from infected tongue shown in **e**, hybridized *in situ* (RNAscope), with probe for MmuPV1 E6 and E7 RNA labeled with diaminobenzidine (DAB). **g,** Section from tongue in **e,f,** showing immunofluorescent (IF) labeling of KRT14 (green) and L1 (pink; not detected). **h,** Lesion severity at indicated time post infection with ∼10^9^ VGE (2-week timepoint data are a subset of data in **c**). WRS test of difference in lesion severity: * p=0.020**. i,** Schematic of T-cell depletion experiment. **j,** Flow cytometry graphs showing CD4+ (upper left) and CD8+ (lower right) populations. Left panel: isotype control antibody treatment; right panel: CD4, CD8 antibody treatment. **k,** Results of T-cell depletion experiment. Two-sided WRS tests of difference in severity: * p=0.038; *** p<10^-7^. Control, 2 weeks vs 4 weeks: p<10^-2^; Depleted, 2 weeks vs 4 weeks: p=0.34.

Remarkably, lesions that developed 2 w.p.i. in 2 mice were diagnosed as squamous cell carcinoma (SCC) by both pathologists (Fig. 1b,c). These cancers developed in mice infected at 8 and 17 weeks of age. Figure 1 shows hematoxylin-and-eosin-stained (H&E) sections of the BoT of a mock-infected control and one of these SCCs (Fig. 1d,e). SCCs expressed MmuPV1 transcripts, assessed by *in situ* hybridization (RNAscope) with probes for MmuPV1 E4 and E6/E7, which identify overlapping sets of MmuPV1 transcripts^25^ (Fig. 1f and Extended Data Table).

Both SCCs expressed the epithelial basal cell marker, Keratin 14 (KRT14) and one expressed the late papillomaviral capsid protein L1 (Fig. 1g; Extended Data Table). Low or absent L1 expression is common among MmuPV1-induced cancers and among HPV-bearing oropharyngeal cancers^26,27,28^. Both cancers were highly proliferative, as assessed by Ki67 expression, and had areas of elevated levels of phosphorylated ribosomal protein S6 (pS6), generally associated with papillomavirus-induced head-and-neck cancers due to activation of the PI3 kinase-mTOR pathway^29^ (Extended Data Fig. 1b,c).

Both SCCs were inflamed and contained koilocytes – histologically abnormal virus-containing cells frequently found in productive papillomavirus-bearing lesions^4,18,28^ (Table). More than half of infected tongues had dysplasia (17/30; data combined from 3 non-contemporaneous experiments). Inflammation and koilocytes were also present in many dysplastic lesions (data combined from 6 experiments; inflammation: 19/57; koilocytes: 51/57; Table). Viral DNA and RNA were detected by *in situ* hybridization (RNAscope) in all neoplastic lesions tested (8/8; Extended Data Table). Mock-infected mice were negative for MmuPV1 as assessed by RNAscope (E6/E7: 0/2; E4: 0/4).

## Base-of-tongue lesions undergo T-cell-mediated regression by 4 weeks

To determine whether neoplastic lesions persist, tissues were collected at 2, 4, and 12 w.p.i. (Fig. 1h; results from 2-week timepoint from this experiment are included in combined results shown in Fig. 1c). No lesions were present at 4 or 12 weeks except for a single case of mild dysplasia at 4 weeks (disease severity at 2 weeks vs 4 weeks, p=0.020). A repeat timecourse yielded similar results (Extended Data Fig. 2b). Non-dysplastic sites of infection at 4 weeks were virtually all inflamed (18/19; Extended Data Fig. 3c).

Previous work demonstrated that BoT lesions in immune-deficient NSG mice infected with MmuPV1 persist to at least 21 weeks^24^. The importance of T cells in preventing MmuPV1-induced benign cutaneous papilloma development and in promoting their regression has been established^19,30,31,32^. To determine whether T cells are also responsible for the eradication of rapid-onset lesions at the BoT, CD4+ and CD8+ T cells were depleted *in vivo* using monoclonal antibodies beginning 4 days prior to infection at the BoT (and anus, discussed below). Tissue was collected 2 and 4 weeks post infection with ∼10^10^ VGE of virus (Prep 3; Fig. 1i). Blood was collected just prior to euthanasia and evaluated by flow cytometry to confirm depletion of CD4+ and CD8+ T cells (Fig 1j).

Depletion of T cells caused rapid-onset lesions to persist. While no mice treated with isotype control antibodies had dysplasia at 4 w.p.i., all mice depleted of T cells had dysplasia at that timepoint (Fig 1k). Notably, T-cell-depleted and control mice differed even at 2 w.p.i.: all depleted mice had dysplastic lesions, whereas 8/19 control mice were lesion free. This significant difference in the frequency of dysplasia (p=0.012) suggests that some or all lesion-free control-treated mice had been infected and developed lesions that were eradicated in a T-cell-dependent manner by 2 w.p.i. Consistent with this hypothesis, lesion-free infected FVB mice treated with control antibodies, assessed 2 w.p.i., were positive for MmuPV1 by RNAscope at the BoT (4/4 tested; Extended Data Table). These lesion-free infected mice were uniformly inflamed (8/8; Table), significantly more than mock-infected mice 2 w.p.i. (1/21, data combined from 5 experiments; p<10^-5^; Table). Similarly, infected, lesion-free mice 2 and 4 w.p.i. in other experiments were significantly more inflamed than mock-infected controls (27/33 vs 1/31; Table; Extended Data Fig. 3b,c). No neoplastic lesions in T-cell-depleted mice were inflamed (Table; Extended Data Fig. 3d). These results indicate that T cells are required for elimination of neoplasia, and that lesion elimination is likely to begin prior to 2 weeks post infection.

**Table.**
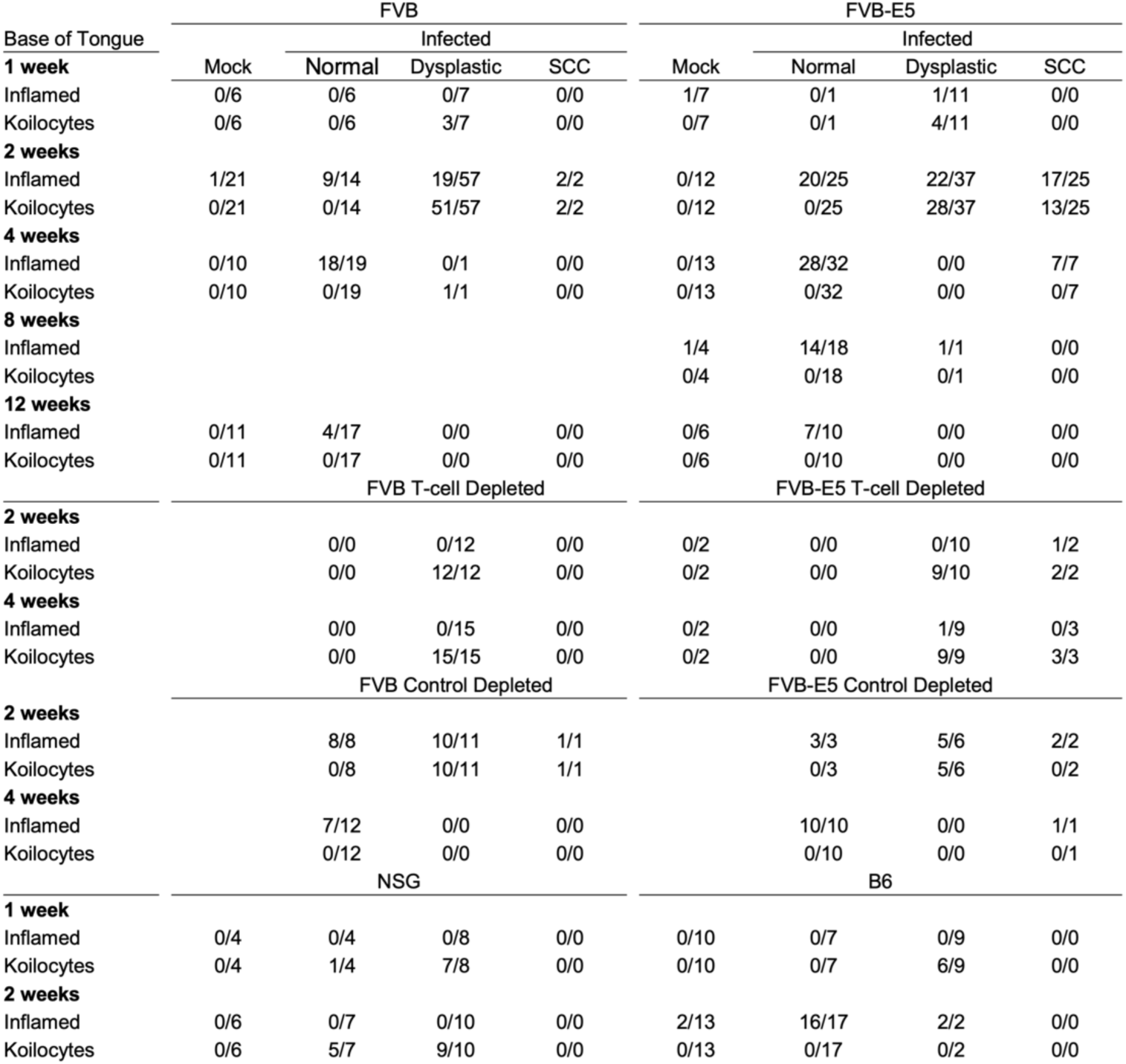
Inflammation and koilocytosis at infected sites (combined data from multiple experiments).

| Base of Tongue | FVB |  |  |  | FVB-E5 |  |  |  |
| --- | --- | --- | --- | --- | --- | --- | --- | --- |
|  | Mock | Infected |  |  | Mock | Infected |  |  |
|  |  | Normal | Dysplastic | SCC |  | Normal | Dysplastic | SCC |
| **1 week** |  |  |  |  |  |  |  |  |
| Inflamed | 0/6 | 0/6 | 0/7 | 0/0 | 1/7 | 0/1 | 1/11 | 0/0 |
| Koilocytes | 0/6 | 0/6 | 3/7 | 0/0 | 0/7 | 0/1 | 4/11 | 0/0 |
| **2 weeks** |  |  |  |  |  |  |  |  |
| Inflamed | 1/21 | 9/14 | 19/57 | 2/2 | 0/12 | 20/25 | 22/37 | 17/25 |
| Koilocytes | 0/21 | 0/14 | 51/57 | 2/2 | 0/12 | 0/25 | 28/37 | 13/25 |
| **4 weeks** |  |  |  |  |  |  |  |  |
| Inflamed | 0/10 | 18/19 | 0/1 | 0/0 | 0/13 | 28/32 | 0/0 | 7/7 |
| Koilocytes | 0/10 | 0/19 | 1/1 | 0/0 | 0/13 | 0/32 | 0/0 | 0/7 |
| **8 weeks** |  |  |  |  |  |  |  |  |
| Inflamed |  |  |  |  | 1/4 | 14/18 | 1/1 | 0/0 |
| Koilocytes |  |  |  |  | 0/4 | 0/18 | 0/1 | 0/0 |
| **12 weeks** |  |  |  |  |  |  |  |  |
| Inflamed | 0/11 | 4/17 | 0/0 | 0/0 | 0/6 | 7/10 | 0/0 | 0/0 |
| Koilocytes | 0/11 | 0/17 | 0/0 | 0/0 | 0/6 | 0/10 | 0/0 | 0/0 |
|  |  | FVB T-cell Depleted |  |  |  | FVB-E5 T-cell Depleted |  |  |
| **2 weeks** |  |  |  |  |  |  |  |  |
| Inflamed |  | 0/0 | 0/12 | 0/0 | 0/2 | 0/0 | 0/10 | 1/2 |
| Koilocytes |  | 0/0 | 12/12 | 0/0 | 0/2 | 0/0 | 9/10 | 2/2 |
| **4 weeks** |  |  |  |  |  |  |  |  |
| Inflamed |  | 0/0 | 0/15 | 0/0 | 0/2 | 0/0 | 1/9 | 0/3 |
| Koilocytes |  | 0/0 | 15/15 | 0/0 | 0/2 | 0/0 | 9/9 | 3/3 |
|  |  | FVB Control Depleted |  |  |  | FVB-E5 Control Depleted |  |  |
| **2 weeks** |  |  |  |  |  |  |  |  |
| Inflamed |  | 8/8 | 10/11 | 1/1 |  | 3/3 | 5/6 | 2/2 |
| Koilocytes |  | 0/8 | 10/11 | 1/1 |  | 0/3 | 5/6 | 0/2 |
| **4 weeks** |  |  |  |  |  |  |  |  |
| Inflamed |  | 7/12 | 0/0 | 0/0 |  | 10/10 | 0/0 | 1/1 |
| Koilocytes |  | 0/12 | 0/0 | 0/0 |  | 0/10 | 0/0 | 0/1 |
|  |  | NSG |  |  |  | B6 |  |  |
| **1 week** |  |  |  |  |  |  |  |  |
| Inflamed | 0/4 | 0/4 | 0/8 | 0/0 | 0/10 | 0/7 | 0/9 | 0/0 |
| Koilocytes | 0/4 | 1/4 | 7/8 | 0/0 | 0/10 | 0/7 | 6/9 | 0/0 |
| **2 weeks** |  |  |  |  |  |  |  |  |
| Inflamed | 0/6 | 0/7 | 0/10 | 0/0 | 2/13 | 16/17 | 2/2 | 0/0 |
| Koilocytes | 0/6 | 5/7 | 9/10 | 0/0 | 0/13 | 0/17 | 0/2 | 0/0 |

## Mice expressing HPV16 E5 develop MmuPV1-induced cancers more frequently

MmuPV1 lacks a homolog of the E5 gene found in high-risk HPVs that cause anogenital and head and neck cancers^33^. HPV16 E5 was shown to have oncogenic properties *in vitro*, as well as *in vivo* using transgenic FVB mice expressing HPV16 E5 in epithelia^34,35,36^. Mice from one of these “FVB-E5” lines ((FVB/NTac-Tg(*KRT14HPV16E5\**)33Plam/Plam)^37^ develop hyperplasia and, with age, mostly benign tumors (6.2% by 15 months, with an average onset of 10.4 months)^34,37^. A previous study showed that infecting FVB-E5 (line 33) mice with MmuPV1 led to earlier detection and more rapid growth of overt MmuPV1-induced lesions in ear skin, as well as increased frequency of SCC at 4 months post infection in the reproductive tract, compared to FVB mice^38^. In addition, less spontaneous ear lesion regression was observed in FVB-E5 mice^38^.

To assess the influence of HPV16 E5 on rapid-onset disease induced by MmuPV1, FVB-E5 (line 33) mice were infected with ∼10^9^ VGE of MmuPV1 (Prep 1) at the BoT, at the same time as non-transgenic FVB mice described above, and lesions at 2 w.p.i. were assessed independently by one or two pathologists blinded to treatment (R.H. and, for a subset, J.P.S.). FVB-E5 mice developed significantly more cancers than non-transgenic mice (Fig. 2a,b). Figure 2 (d,e) shows H&E-stained sections of a mock-infected control and one of the SCCs found in an infected FVB-E5 mouse. As in FVB mice, cancers in FVB-E5 mice expressed MmuPV1 transcripts; expressed KRT14; were highly proliferative; and had areas of elevated pS6 expression (Fig. 2f,g; Extended Data Table; Extended Data Fig. 1b). Cancers arising in infected FVB-E5 expressed L1, but only in a few cells (Fig. 2g; Extended Data Table).

**Figure 2.**
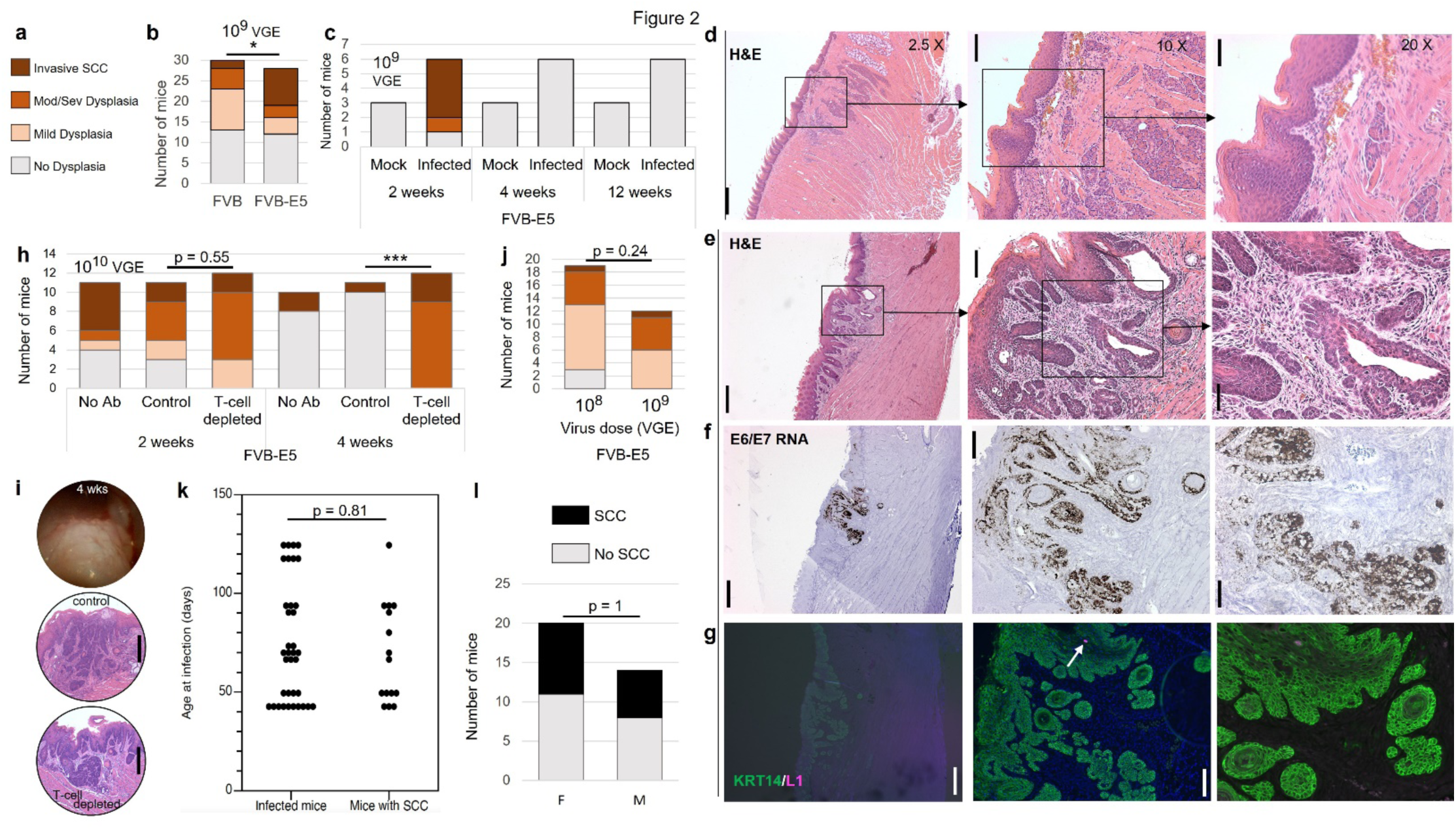
FVB-E5-transgenic mice develop more rapid-onset oropharyngeal SCCs than non-transgenic FVB mice. **a,** Legend for graphs in **b**,**c**,**h**,**j**. **b,** Lesion severity in FVB and FVB-E5 (“E5”) mice infected with ∼10^9^ VGE MmuPV1 at BoT. Fisher’s Exact (FE) test of difference in cancer frequency, 3 experiments combined: * p=0.019. WRS test of difference in overall lesion severity p=0.27. **c,** Lesion severity of FVB-E5 mice at indicated time post infection with ∼10^9^ VGE. **d,** Base of FVB-E5 tongue mock-infected with PBS, stained with H&E. Panels are images of a single section from one mouse. Scale bars: 400 um (2.5X objective), 100 um (10X objective), 50 um (20X objective). **e,** SCC at base of FVB-E5 tongue infected with 10^9^ VGE of MmuPV1, stained with H&E. Panels and scale bars as in **d**. **f,** Section from infected tongue shown in **e**, hybridized in situ (RNAscope), with probe for MmuPV1 E6 and E7 RNA labeled with DAB. **g,** Section from tongue in **e, f,** showing IF labeling of KRT14 (green) and L1 (pink; arrow points to single positive cell detected). **h,** T-cell-depletion experiment performed as in Fig. 1, with additional “No antibody” groups. Two-sided WRS tests of severity, control vs depleted at 4 weeks: *** p<10^-3^. Disease severity, 2 weeks vs 4 weeks: No Ab, p=0.086; Control, p<10^-2^; Depleted, p=0.030. **i,** Endoscopic image (top) and H&E-stained section (middle) of SCC from same 4 w.p.i. control-antibody-treated mouse; bottom: H&E-stained section of SCC from 4 w.p.i. T-cell-depleted mouse. Scale bar=400 um. **j,** Lesion severity in FVB-E5 mice infected with 10^8^ vs 10^9^ VGE at BoT; difference assessed by WRS. **k,** Graph of the age of individual mice, represented by a filled circle, at infection (left column) vs the subset of those infected mice that developed SCC (right column); difference assessed by WRS. **l,** Cancer frequency in female vs. male FVB-E5 mice; difference assessed by FE test.

## Cancers in FVB-E5 mice regress

The higher frequency of SCC in FVB-E5 mice allowed us to address whether these cancers regress. Whereas 4/6 mice developed SCC at 2 weeks post infection with ∼10^9^ VGE (Prep 1), no neoplastic lesions were detected in tongues collected 4 or 12 w.p.i. (Fig. 2c). This reduction in SCC frequency was significant (2 weeks vs 4 and 12 weeks combined: p<10^-3^), indicating these rapid-onset cancers regress. A repeat of this time course with a lower dose of virus (∼5 x 10^8^ VGE; Prep 2) also showed complete regression of all neoplastic lesions (Extended Data Fig. 2e). Extended Data Figure 2 shows CD4+ and CD8+ cells in an oropharyngeal SCC at the BoT 2 w.p.i. (Extended Data Fig. 2f).

*In vivo* T-cell depletion was performed to determine whether tumor regression was mediated by T cells. At 4 weeks post infection with ∼10^10^ VGE at the BoT, all T-cell-depleted mice (12/12) had dysplastic lesions and SCCs, whereas 18/21 control- or no-antibody-treated mice were lesion free (Fig. 2h). Lesion-free control-antibody-treated FVB-E5 tongues at 2 and 4 w.p.i. were inflamed at the inoculation site (3/3, 10/10; Table), while mock-infected tongues 2 w.p.i. were never inflamed (0/12; results combined from 4 experiments; Table) and the neoplastic lesions of T-cell depleted mice were rarely inflamed (2/24; Table; Extended Data Fig. 3d). Infected, lesion-free mice 2 and 4 w.p.i. in other experiments were also significantly more inflamed than mock-infected controls (48/57 vs 0/25; Table; Extended Data Fig. 3b,c). These results demonstrate that T cells mediate oropharyngeal lesion regression in infected FVB-E5 mice as observed in non-transgenic FVB mice (Fig. 1). The presence of inflammation at 2 weeks in infected FVB-E5 mice that had no observable lesions suggests that regression began prior to 2 w.p.i. in these mice as in FVB mice.

The few control/no antibody mice with disease remaining at 4 w.p.i. exclusively had SCC (3/21; Fig. 2h). Figure 2i shows an endoscopic image of one of these cancers. An H&E-stained section of that cancer (center panel) shows inflammation consisting predominantly of lymphocytic infiltrate with neutrophils. An uninflamed T-cell-depleted cancer at 4 w.p.i is shown in the bottom panel (higher magnification of H&E panels: Extended Data Fig 3e,f). To determine whether cancers are present at later time points in FVB-E5 mice infected with this high dose of ∼10^10^ VGE (Prep 3), tongues were collected 4 and 8 w.p.i. SCCs were found in 5/15 of mice at 4 weeks, but none were detected at 8 weeks (p=0.011; Extended Data Fig. 2g), indicating that cancers also regressed after infection with this dose of virus.

SCCs in T-cell-depleted mice at 4 weeks had koilocytes and lacked inflammation (koilocytes: 3/3; inflammation: 0/3). In contrast, cancers in control- and no-antibody-treated mice at 4 weeks uniformly lacked koilocytes and were inflamed (0/8, 8/8; results combined from 2 experiments; Table; Extended Data Fig. 3e,f). The lack of koilocytes in control mice at 4 weeks represents a significant loss relative to 2 w.p.i. Differences in koilocytosis and inflammation between control and T-cell-depleted cancers at 4 w.p.i. are significant (Extended Data Fig. 3g,h,i).

Cancer frequency at the BoT in FVB-E5 mice was not affected by viral dose, in a comparison of 10^8^ and 10^9^ VGE/ul diluted from the same virus stock (Fig. 2j); age at infection (6-18 weeks; Fig. 2k); sex (Fig. 2l); or filtration of virus through a 0.23 um filter to eliminate contaminating cells and particles larger than viruses (Prep 3; Extended Data Fig. 2h). Viral DNA and/or RNA was detected in paraffin sections of 18/21 tested FVB and FVB-E5 BoT SCCs (Extended Data Table). No metastases were detected in lungs and regional lymph nodes collected 4 w.p.i. in control- or antibody-treated mice.

## Base-of-tongue lesions develop within one week and persist in some strains

Moderate to severe dysplasia can develop within 7-14 days of infection in the reproductive tract in B6 (C57BL/6J) mice and in the larynx, trachea, and palate of immune-deficient NSG mice (NOD.Cg-*Prkdc^scid^-Il2rg ^tm1Wjl^/*SzJ)^21,22^. To determine when oropharyngeal dysplasia develops, FVB, FVB-E5, B6, and NSG mice were infected at the BoT and collected 1 w.p.i. Papillomavirus-induced dysplasia was present at the BoT in the majority of mice in all strains tested (Extended Data Fig 4). These results are similar to those reported by Scagnolari et al. for MmuPV1-induced vaginal lesions in B6 at 7-11 days post infection, except for severity: these BoT lesions in B6 were all mildly dysplastic, whereas vaginal lesions were moderately to severely dysplastic^21^.

Little to no inflammation was observed histologically at 1 week in FVB, FVB-E5, or B6 mice (2/64; Table), in contrast to frequent inflammation in infected tissues at 2 weeks (above; Table). NSG mice developed no inflammation at either timepoint (1 week: 0/16; 2 weeks: 0/23; Table).

Few B6 BoT lesions remained at 2 weeks (2/15), consistent with the elimination of vaginal lesions in B6 mice between 11 and 30 days^21^. One quarter (2/8) of non-dysplastic, infected B6 mice were positive for viral DNA/RNA 2 w.p.i.; however, these infection sites were weakly labeled (Extended Data Table, Extended Data Fig. 4d). In contrast to BoT lesions in B6, lesions in FVB, FVB-E5, and NSG mice included moderate-to-severely dysplastic lesions at 1 week as well as at 2 w.p.i. One FVB lesion at 1 w.p.i. was scored as at least severe dysplasia with foci suspicious for early invasive carcinoma (Extended Data Fig. 4c). The persistence of disease in NSG mice is consistent with prior observations of severe dysplastic disease but not cancer at 21 weeks in the oropharynx^24^.

## The oral tongue is resistant to rapid-onset cancer

HPV-associated cancer is far more prevalent in the oropharynx than the oral cavity^23,39^. To determine whether the oropharyngeal BoT is more susceptible to rapid-onset papillomavirus-induced carcinogenesis than the oral, anterior, tongue, FVB-E5 mice were infected at each site separately (Fig. 3a). Lesions were assessed by a pathologist blinded to treatment (R.H.).

**Figure 3.**
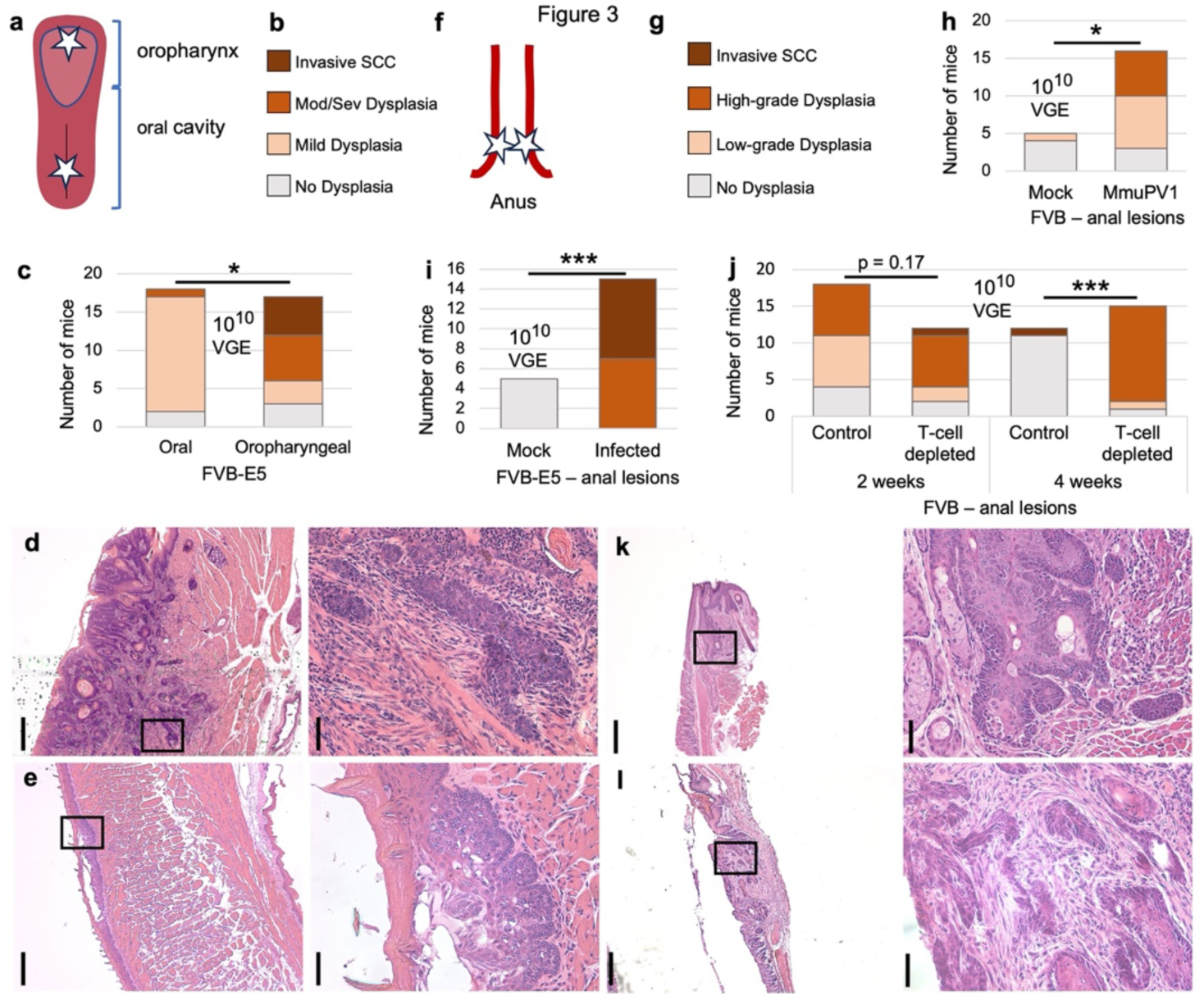
The anus, but not the oral tongue, is susceptible to rapid-onset cancer and T-cell-mediated regression. **a,** Illustration of the oropharyngeal (base of) tongue, and the oral (anterior) tongue, with stars showing the respective sites of infection. **b,** Legend for graph in **c**. **c,** Lesion severity in mice infected with 10^10^ VGE MmuPV1 using the Greer Pick in the oral or oropharyngeal tongue. FE test of difference in cancer frequency, * p=0.019; WRS test of difference in overall lesion severity, p<10^-2^. **d,e,** Scale bars in left panels: 400 um; right panels: 50 um. **d,** FVB-E5 invasive oropharyngeal (BoT) cancer 2 w.p.i. **e,** FVB-E5, moderately dysplastic oral tongue lesion 2 w.p.i. **f,** Illustration of the lower gastrointestinal tract with stars showing the sites of infection in the anus. **g,** Legend shows the categories of dysplasia used in assessing anal disease in **h-j**. **h,** Lesion severity in FVB mice infected with 10^10^ VGE or mock-infected with a corresponding non-viral control skin prep. WRS test of difference in lesion severity: * p=0.014. **i,** Lesion severity in FVB-E5 mice infected with 10^10^ VGE or mock-infected with control skin prep. WRS test of difference in severity: *** p<10^-3^. **j,** T-cell depletion experiment as in Figure 1i. Animals were infected both at the base of the tongue, shown in Figure 1, and at two locations in the anus with 10^10^ VGE. WRS test of difference in lesion severity in the anus at 4 w.p.i: *** p<10^-4^; Control 2 weeks vs 4 weeks p<10^-2^; T-cell depleted 2 weeks vs. 4 weeks p=0.49. **k,l,** Scale bars in left panels: 400 um; right panels: 50 um. **k,** FVB-E5, anal SCC 2 w.p.i. **l,** T-cell-depleted FVB, anal SCC 2 w.p.i.

While most of the mice infected at the BoT developed SCC (5/17) or moderate-to-severe dysplasia (6/17), mice infected in the anterior tongue developed no cancers, few moderately dysplastic lesions (3/32), and many mildly dysplastic lesions (26/32; 2 combined experiments; Fig. 3c-e and Extended data Fig 2i). The difference in cancer frequency is significant (Fig. 3c). Injury and infection at one site did not lead to secondary infection at the other site in the same animal, within this short timeframe. These results indicate that, although the oral tongue can be infected and develop dysplastic lesions within two weeks of infection, it is resistant to rapid-onset SCC.

## The anus, like the oropharynx, is susceptible to rapid-onset cancer and T-cell-mediated lesion regression

Like the oropharynx, the anus is susceptible to human papillomavirus-induced cancer^40^. MmuPV1 can synergize with a chemical carcinogen to induce anal cancer within 6 months in immune-deficient mice^41^. To assess whether MmuPV1 can induce rapid-onset cancer in immune-competent mice, FVB and FVB-E5 mice were infected with ∼10^10^ VGE at two sites in the anus, 180° apart, using the Greer Pick (Fig. 3f). Lesions were assessed by a pathologist blinded to treatment conditions (K.A.M.). Among FVB mice, 2 weeks after infection with 10^10^ VGE, 13/16 developed dysplastic lesions, with 6/16 severely dysplastic (Fig. 3h). Among FVB-E5 mice, all developed neoplastic lesions, and half of these (8/15) were SCCs (Fig. 3i,k).

MmuPV1 infection of the anus in immune-deficient NSG mice persists and causes severely dysplastic lesions by 6 months post-infection^41^. In contrast, FVB mice infected anally do not have significantly more dysplastic anal tissue than mock-infected controls at 6 months. To determine whether anal lesions present 2 w.p.i. undergo T-cell-mediated regression, as do BoT lesions, the anus of T-cell-depleted FVB mice was infected (as well as the BoT; see Figure 1i,j). There was no difference in penetrance/severity of anal disease in the T-cell-depleted versus control (isotype-antibody-treated) mice at 2 w.p.i., although one T-cell-depleted mouse developed invasive SCC (Fig. 3l). By 4 w.p.i. the difference between treatments was significant, with 14/15 T-cell-depleted mice having neoplastic disease compared to only 1/11 control-treated mice (Fig. 3j). The reduction in disease severity between 2 and 4 weeks for control mice was also significant (p<10^-2^). As in the FVB-E5 oropharyngeal T-cell-depletion experiment (Fig. 2h), the sole remaining dysplastic lesion in control-treated FVB mice at 4 weeks was a SCC.

## The skin is susceptible to rapid-onset cancer and lesions that persist

The mouse papillomavirus MmuPV1 was first discovered as a virus that caused cutaneous warts in nude mice^18^. Infection was shown to cause invasive cancers in ear skin by 16 w.p.i. in immune-competent mice^42^. To determine whether invasive cancer can arise within 2 weeks of infection in ear skin, the ventral face of each ear (Fig 4a) was scarified with a needle, and 2 ul of virus was applied to the wound (10^9^ or 10^10^ VGE, Prep 3; Fig. 4a). Scoring of these lesions includes an additional category - “possibly invasive” (Fig. 4b) - because in some cases pathologists (R.H., J.P.S, and D.B., assessing overlapping sets of experiments) could not determine whether islands of dysplastic cells were growing along epithelial appendages (e.g. sebaceous glands).

**Figure 4.**
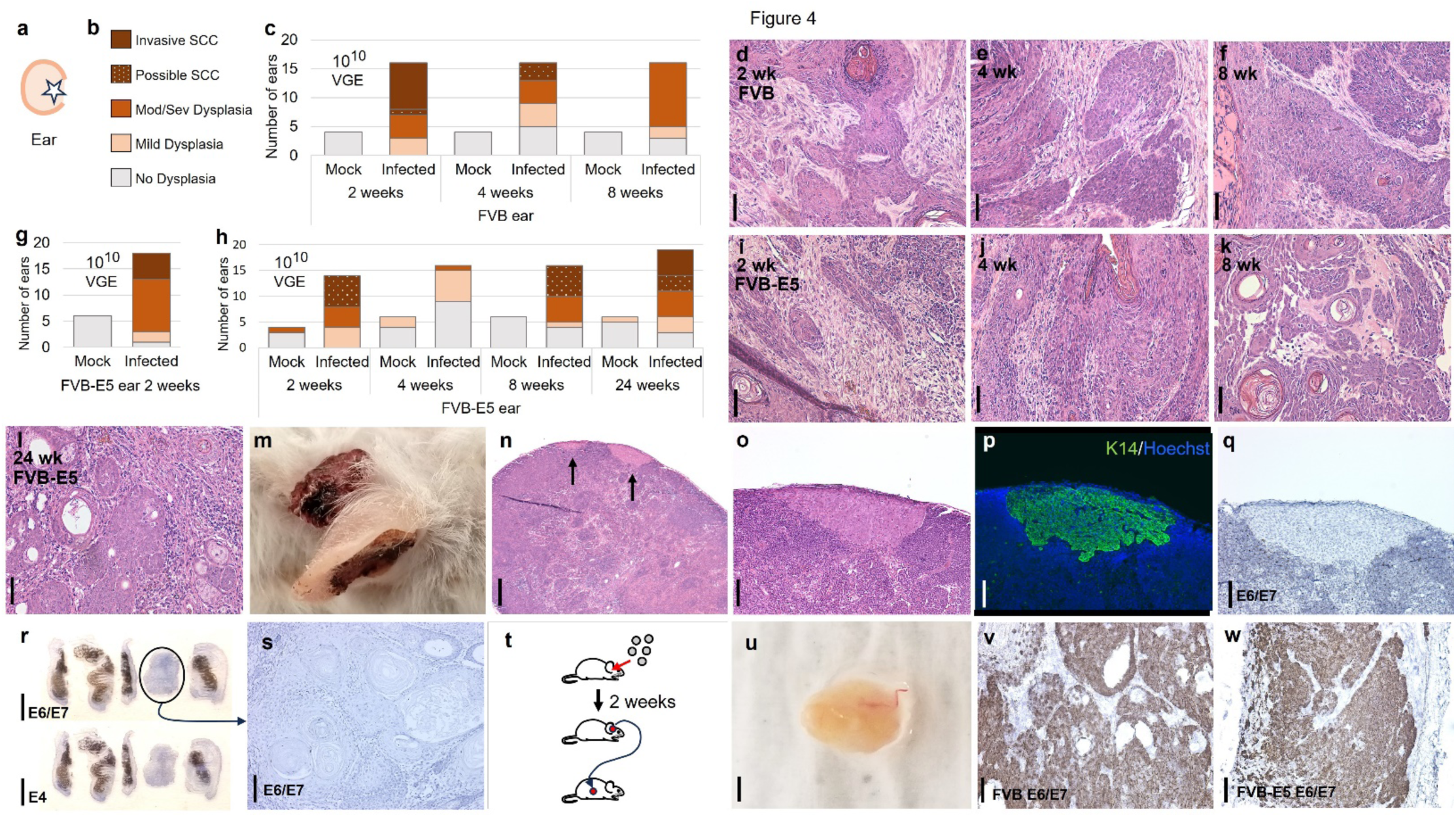
The skin is susceptible to acute cancer and disease that persists. **a,** Illustration of ear and location of scarification and infection on the inner surface of the ear, indicated by a star. **b,** Legend for disease severity in **c,g,h**. **c,** Severity of FVB lesions at indicated time post infection of each ear with ∼10^10^ VGE MmuPV1 or mock-infection with non-viral control skin prep. **d-f,** FVB ear lesions from **c** stained with H&E. Scale bar=50 um. **d,** SCC 2 w.p.i. **e,** Possible SCC 4 w.p.i. **f,** Severe dysplasia 8 w.p.i. **g,** Severity of FVB-E5 lesions collected two weeks post infection of each ear with ∼10^10^ VGE MmuPV1 or mock infection with control skin prep. **h,** Severity of FVB-E5 lesions, in a separate experiment, collected at indicated timepoints post infection with ∼10^10^ VGE or mock infection with control skin prep. One ear (24 weeks) showed atypia that was difficult to categorize and was therefore left out of this analysis. **i-l,** Sections of FVB-E5 ear lesions from **h** stained with H&E. Scale bar=50 um. **i,** Possible SCC 2 w.p.i. **j,** Severe dysplasia 4 w.p.i. **k,** Possible SCC 8 w.p.i. **l,** SCC 24 w.p.i. **m,** Right ear lesion in FVB-E5 mouse 38 w.p.i. with metastasis to lymph node. **n,** H&E-stained section of lymph node metastasis from mouse in **m**. Scale bar=400 um. **o-q,** Serial sections of lymph node with metastasis. Scale bar=100 um. **o,** Stained with H&E. **p,** Labeled with antibody to KRT14 and Hoechst dye. **q,** labeled with probes for MmuPV1 E6/E7 using RNAscope. **r,** Ear sections from mouse with metastasis shown in **n**, labeled using RNAscope with probes for MmuPV1DNA/RNA. Top panel: E6/E7 probe. Bottom panel: E4 probe. Scale bar=5 mm. **s,** Section of invasive ear lesion in Fig. 4r that had no detectable MmuPV1 E6/E7 DNA/RNA (circled). Scale bar=50 um. **t,** Illustration of grafting procedure. FVB and FVB-E5 ears were infected with ∼10^10^ VGE MmuPV1. After 2 weeks, lesions were excised and divided into fragments that were implanted subcutaneously and allowed to grow ∼4 months. **u,v,** FVB ear lesion graft 17 weeks after subcutaneous implantation. **u,** whole cystic oval graft. Scale bar=1 mm. **v,** section of **u** labeled with probes for MmuPV1 E6/E7 using RNAscope. Scale bar=50 um; **w,** FVB-E5 graft 16 weeks after implantation; labeled as for **v**. Scale bar=50 um.

Infection of either FVB or FVB-E5 mice caused SCCs to develop by 2 w.p.i., and 95% of ears had neoplastic lesions at this timepoint (80/84; Fig. 4c,d,g-i and Extended Data Fig. 5a,d,m-n). Notably, lesions persisted to 8 weeks in both strains (Fig. 4c-k; Extended Data Fig. 5a-f,m,n); although disease severity decreased significantly (chi-square, combined doses: FVB, p<10^-5^; FVB-E5, p<10^-5^). Ears were inflamed at each timepoint (FVB 10^10^ VGE timecourse, Fig. 4c: 2 weeks, 16/16; 4 weeks, 13/16; 8 weeks, 14/16; mock-infected: 0/6). FVB-E5 were also assessed for lesions at 24 w.p.i (Fig. 4h,l and Extended Data Fig. 5g,o). At this timepoint SCCs were detected using both doses of virus, consistent with previous observations in FVB and FVB-E5 mice 4-6 months post infection with 10^8^ or 10^10^ VGE^38,42^. In FVB mice, while disease severity decreased with time at each dose, overall disease severity was higher after infection with the higher dose (chi-square: p<10^-3^) consistent with previous observations^38,43^.

Combining results for both strains and virus dosages, a pattern emerged: approximately one-third of lesions regressed and the frequency of frank invasive SCC decreased significantly between 2 and 4 weeks (lesion frequency: 95% vs 62%; p<10^-6^; SCC frequency:18% vs 1.6%, p<10^-2^). While the frequency and overall severity of lesions did not change significantly between 4 and 24 weeks (frequency: 4 vs 8 weeks: 62% vs 67%, p=0.59; 8 vs 24 weeks: 67% vs 63%, p=0.83; severity: 4 vs 8 weeks, p=0.21; 8 vs 24 weeks, p=0.78), the frequency of cancer increased again, significantly, between 8 and 24 weeks (1.6% vs 9.4%, p=0.021). These results indicate that both FVB and FVB-E5 mice can develop invasive SCC of ear skin within 2 weeks of infection; that, in contrast to the oropharynx and the anus, dysplastic disease largely persists; and that cancer frequency ebbs between 2 and 24 weeks.

## MmuPV1-induced cancer can metastasize

At 38 weeks post infection of both ears, the ears, lungs, and cervical lymph nodes were collected from a mouse with large ear lesions, one of which extended into the skin at the back of the head (Fig. 4m and Extended Data Fig 5h). The ear lesions were diagnosed as SCCs, with metastatic foci found in one cervical lymph node (Fig. 4n-q; Extended Data Fig. 5i). Metastatic cells were positive for KRT14 (Fig. 4p) and TRP63 (not shown) but viral DNA was not detectable (Fig. 4q and Extended Data Fig. 5i). Notably, a portion of one of the same mouse’s ear lesions also failed to label positively for MmuPV1 DNA/RNA with probes for either E6/E7 or E4 (Fig. 4r,s and Extended Data Fig 5j,k). This portion was also scored as SCC and was positive for KRT14 and TRP63 (Extended Data Fig. 5l) and was therefore a likely primary source of the metastatic cells. Lack of detectable viral DNA specifically associated with invasive portions of MmuPV1-induced tumors has been reported^44^. No lung lesions were detected.

## Lesions that develop by 2 weeks can grow independently as grafts

A classic test for malignancy involves ectopic (e.g. subcutaneous) implantation of the putative invasive cancer into a new host mouse^45,46^. Malignant human and mouse cells continue to grow under these conditions, while non-malignant cells – even rapidly growing embryonic cells – do not^46^. To determine whether MmuPV1-induced lesions 2 w.p.i. can grow independently as ectopic tumor grafts, 20 FVB and 20 FVB-E5 ears were infected with ∼10^10^ VGE MmuPV1. Two weeks later, pieces of the 8 largest FVB lesions and the 10 largest FVB-E5 lesions were grafted subcutaneously in adult NSG mice (Fig. 4t). Lesions were measured weekly. Two lesions grew quickly and were collected when they reached approximately 100 mm^3^: one FVB lesion, collected ∼17 weeks post transplantation and one FVB-E5 lesion collected at 16 weeks (Extended Data Fig. 6a). Both grafts had transformed into oval cysts, as commonly seen with HPV+ cancer grafts (Fig. 4u)^47^. Sections of these grafts revealed extensive tumor tissue (Extended Data Fig 6b,c) positive for MmuPV1 DNA/RNA by RNAscope using E6/E7 probes (Fig. 4v,w). These results provide independent evidence corroborating pathologists’ observations that MmuPV1-induced lesions can become invasive within 2 weeks of infection.

## Discussion

To our knowledge, the development of squamous cell carcinoma within 2 weeks of papillomavirus infection is the most rapid onset of invasive cancer reported in any animal. The fact that these epithelial cancers can be induced with a naturally occurring virus in immune-competent, wild-type, adult animals, in the absence of carcinogenic cofactors, is even more remarkable. Most previously reported examples of rapid invasive cancer development in people and animal models have involved the specialized environment of developing tissues, combinations of carcinogenic elements such as transgenic oncogenes and mutagens, or immune suppression. Examples include: congenital tumor development followed by invasive cancer detection in the first weeks after birth (e.g. in human infant neuroblastoma, where 16% are diagnosed during the first month following birth^48^; in the Mouse Mammary Tumor Virus - Polyoma Middle T-(MMTV-PyMT) antigen model^49^; or in the Melanoma-Bearing Libechov Minipig pig^16^); viral infection of newborns followed by invasive cancer detection approximately 2 months later (e.g. Maloney Murine Sarcoma Virus in rats^50^); induction of oncogene expression together with weeks of exposure to carcinogen leading to invasive cancer within 9 weeks^51^; the development of EBV-induced lymphoma within 2 months of organ transplant in chemically immune-suppressed patients ^6–8^; intraperitoneal injection of EBV-infected human cord blood cells leading to invasive lymphomas in immune-deficient mice within 4 weeks^9^; and the simultaneous induction of multiple transgenic oncogenes together with mutation of a tumor suppressor leading to invasive cancer within 20 days^52^. By contrast, the cancers we observe 2 weeks post infection are caused by papillomavirus alone, in adult mice that have not been mutated or treated with carcinogens and that have intact immune systems.

In some previously described cases of rapid-onset cancer, visible growths were not analyzed at earlier time points, or authors did not specify when invasive tumors were first detected; cancers might therefore have developed earlier than reported. Indeed, it seems likely that our observation that SCCs can arise acutely is not unique – that other cancers arise rapidly but fail to be detected or analyzed histologically until they have enlarged beyond a threshold size that is reached months or years after the cancer’s establishment. At two weeks post infection, MmuPV1-induced oropharyngeal and anal SCCs in our experiments were not obvious to the naked eye. Similarly, human cancers at the BoT are often too small to detect using standard methods^11^. Recent modeling of HPV+ head-and-neck cancer development indicates that important cancer drivers such as HPV integration and PIK3CA amplification occur 20-30 years before diagnosis^53^. Indeed, recent analysis of circulating tumor DNA (ctDNA) in patients with oropharyngeal SCC revealed that ctDNA sharing the cancer’s mutational signature was found in patients’ blood samples, collected prospectively, up to 10 years before the cancer was diagnosed^54^ – indicating that human SCC can develop much more rapidly than previously thought.

FVB (FVB/N) mice are not particularly susceptible to spontaneous squamous cell tumors^55^. They are, however, unusually susceptible to SCCs induced by carcinogens, transgenically expressed H-ras, or MMTV-PyMT^56,57,58^; and FVB keratinocytes are more susceptible than those of other strains to malignancy following immortalization by H-ras expression or HPV E6/E7^59^. A polymorphism between the FVB and B6 (C57BL/6) strains, within the *Patched* gene of the Sonic Hedgehog pathway, has been shown to control susceptibility to H-ras-initiated SCCs^56^. The genetic basis for the greater susceptibility of FVB mice than B6 to MmuPV1-induced tumorigenesis^19,43^ (Extended Fig. 4) is not known.

A priori, oncogenic viruses seem more likely than carcinogens or DNA replication errors to induce cancer acutely in adult mice. Viruses such as HPV16 express proteins that can activate oncogenes, inactivate tumor suppressors, induce aneuploidy and chromosome instability, and affect other hallmarks of cancer^1,60^. In contrast, the odds of simultaneously mutating or epigenetically modifying multiple host genes necessary for cancer development, while maintaining cell viability, are likely low. Some experimental support for this hypothesis comes from studies of the rodent liver, which is particularly susceptible to chemical carcinogenesis according to the National Toxicology Program’s two-year bioassay^61^. Injection with a single dose of the carcinogen diethylnitrosamine near its Lethal Dose 50 (dose given: 150-175 mg/kg; LD50: ∼200 mg/kg) caused no detectable hepatocellular carcinomas until 447 days after injection^61^.

If cancers that develop within weeks due to viral infection are a more general phenomenon, our data suggest that many that arise might regress before they can be diagnosed. In our studies, MmuPV1-induced lesions in the oropharynx and anus, including SCCs, regressed completely. T cells were required for disease regression. T cells are also implicated in the regression of melanomas: they extensively infiltrate and become predominant during regression of these cancers in swine models and human patients^62^. The regression of invasive lesions in MmuPV1 -infected tissues represents the first genetically tractable model of spontaneous cancer regression that does not involve grafted cancer cells.

Our results suggest koilocyte loss might be an early response to T cells in papillomavirus-induced cancers. Koilocytes -- abnormal, virus-containing cells -- were present in half of base-of-tongue SCCs at 2 weeks but were absent from SCCs remaining at 4 weeks unless T cells were depleted. T-cell mediated regression might lead to complete elimination of virus-infected cells, or it might result in cells with virus that is subdued: not replicating or replicating at low levels (“latent”^63^), or integrated and not capable of replication. Virus-infected cells remaining after lesion regression could, in theory, continue to grow in response to immune deficiency^63^ or after acquiring new mutations. These possibilities each have clinical implications and warrant careful molecular analysis of regressed lesions.

Lesion persistence differed dramatically between the base of tongue or anus and the ear epidermis. While infected mice had few lesions in the tongue or anus at 4 weeks and none in the tongue at 12 weeks post infection, most mice had lesions in the ear at 4, 8, and 24 weeks post infection. In the ear, the frequency of SCCs declined between 2 and 4 weeks, only to rebound again between 8 and 24 weeks, consistent with previous studies showing the presence of cutaneous ear SCC in both FVB and FVB-E5 mice 4-6 months post infection^38,42^. A mouse with an ear lesion that persisted for 9 months developed lymph node metastases (the first reported for this virus, to our knowledge). While invasive base-of-tongue SCCs were observed in non-transgenic FVB mice 2 weeks post infection, these cancers were rare: only 3 of 84 mice developed SCC. A significantly larger fraction of ears in FVB mice, 8 of 35, developed invasive SCC by 2 weeks, suggesting that the ear is more susceptible to establishment of invasive cancer as well as to persistent disease. MmuPV1 was originally identified as a cutaneous virus and might have evolved to optimize its persistence in that tissue. Spatial molecular analysis could help determine whether partial regression, recurrence, and metastasis correlate with changes intrinsic or extrinsic to the cancer cell.

All FVB ear SCCs present at 2 weeks post infection developed after infection with the higher of two doses tested (10^10^ VGE). Ears infected with the lower dose were first diagnosed with cancer at 4 weeks. This dose dependence of SCC induction in FVB was significant (p<10^-3^) and is consistent with the known dose dependence of the speed and frequency with which MmuPV1 induces benign papillomas in the skin^43^. Higher doses of virus are likely to infect more cells, which in turn could lead to papilloma or SCC development through the cooperation of separately infected cells in a chimeric tumor^63^, or through an increased chance of stochastic carcinogenic changes intrinsic to the cell, such as chromosomal instability-induced aneuploidy, chromothripsis, or the formation of oncogenic cellular-viral extrachromosomal circular DNA hybrids^1,64,65^.

The HPV16 E5 transgene increased MmuPV1-induced severe disease/cancer frequency in the oropharynx, anus, and skin. Proposed mechanisms of E5 oncogenicity include the stimulation of proliferation by upregulation of Epidermal Growth Factor Receptor (EGFR) and other growth signaling pathways, and interference with antigen presentation by tumor cells leading to immune suppression^1,36^. In the oropharynx, transgenic E5 did not confer resistance to spontaneous regression or significantly reduce inflammation. Mice with a dominant-negative mutation in EGFR have been used to show the receptor’s involvement in hyperplasia induced by E5^37^ and can be used to determine EGFR’s involvement in acute onset of invasive cancer. In addition, molecular analyses should point to critical factors modulated by E5.

While many SCCs developed in response to MmuPV1 infection of FVB-E5 mice at the oropharyngeal base of tongue, none developed in the oral (anterior) portion of the tongue. This tissue specificity within the tongue echoes that seen in people and indicates that the presence of a lingual tonsil, which mice do not have, is not required for SCC susceptibility at the base. Circumvallate papillae are associated with the base of the tongue in both species and have previously been shown to be particularly susceptible to mouse papillomavirus infection, although the biological basis for this susceptibility has not been established^66,67^. The susceptible cells at the base of tongue in mice could be the pluripotent cells at the squamous-columnar junction of the salivary glands and the surface epithelium, which reside at the base of the circumvallate papilla and surrounding tissue in mice^68^ (Fig. 1e; Fig. 2e). Such squamous-columnar transition zones are consistently found at tissue sites most susceptible to papillomavirus-induced SCC, including the cervix, anus, and hair follicles of the skin^17,42^.

Our findings, together with recent results of HPV ctDNA analysis^54^, raise the possibility that invasive cancers might arise and regress rapidly in people. If our results translate to humans with HPV infections, acute onset and regression are likely to be rare and tissue specific. More studies are needed to determine the set of molecular and cellular factors that drive these rapid changes and determine which lesions persist and metastasize. Given the speed of these events, progress should be swift.

## Materials and Methods

### Mice

Animal experiments were approved July 3, 2020 and June 14, 2023 by the School of Medicine and Public Health Animal Care and Use Committee (IACUC) of the University of Wisconsin-Madison and conducted in accordance with the National Institutes of Health Guide for the Care and Use of Laboratory Animals (Protocol M005871). Immune-competent mice (FVB/NTac (Taconic Biosciences), FVB-E5 (FVB/NTac-Tg(KRT14-HPV16E5*) 33Plam/Plam), and B6 (C57BL/6J; JAX, stock #000664)) were fed Teklad 2019 diet (Inotiv), while immune-deficient nude mice(Fox1^nu^; Envigo) and NSG mice (NOD.Cg-*Prkdc^scid^-Il2rg ^tm1Wjl^*/SzJ; JAX, stock #005557), purchased from the Jackson Laboratory and bred in a breeding core (BRMS, UW-Madison), were fed irradiated Teklad 2919 diet (Inotiv).

### Virus preparation/control skin preparation

Virus stocks were prepared as described^24^. Briefly, warts were collected from infected areas on the ear and tail of nude mice, as well as from secondary warts that developed elsewhere on the skin. Warts were homogenized, treated with Triton X-100, collagenase, and benzonase followed by centrifugation, treatment with additional benzonase, and collection of the supernatant. “Control skin prep” was prepared in parallel, identically, except instead of warts, skin was collected from uninfected nude mice in locations that corresponded to the location of warts used for virus preparation. The viral genome equivalents present in each virus stock were determined by comparison to DNA standards. Concentrations (VGE/ul) were as follows: Prep 1, ∼ 3 x 10^9^; Prep 2, ∼ 3 x 10^8^; Prep 3, ∼8 x 10^9^.

### Infection

Infected mice were male and female, 4-21 weeks old. Groups within an experiment were populated with mice of a similar range of ages. Mice were infected as follows:

#### Oropharynx/base of tongue and oral cavity/anterior tongue

Mice were anesthetized using isoflurane (Midwest Veterinary Supply or Dechra) provided via Mickey’s Space Helmet^24^ (available from MediLumine). A Greer Pick (Stallergenes-Greer, London, UK) was used to infect the BoT as described^24^. Briefly, the Pick was plunged into virus stock, control skin prep, PBS, or Evans Blue (Sigma) and cleaned of excess liquid by sliding the Pick along the side of the container, leaving ∼1.5 ul in the tip of the Pick. The Pick was then inserted through the oral cavity, turned toward the dorsal surface of the BoT just beyond the visible tongue, pressed into the tongue, and rotated approximately one quarter turn (Extended Data Fig.1a). Infection of the oral/anterior tongue was done in the same way, except the Pick was pressed into the dorsal surface of the anterior tongue approximately 3-5 mm from the tip of the tongue.

#### Anus

Mice were anesthetized using isoflurane and a standard nose cone. A Greer Pick was prepared as above and pressed into the anal canal approximately 2-4 mm from the exterior. A second Pick loaded with virus was pressed into the anal canal at 180° from the first infection site.

#### Skin

Mice were anesthetized using isoflurane and a standard nose cone. An approximately 20 mm^2^ patch on the ventral face of each ear was scarified with a 27-gauge needle, and 2 ul of virus stock was applied to the wound.

### Grafts

Ear lesions were collected from FVB or FVB-E5 mice 2 w.p.i. as follows. In a biosafety cabinet, a piece of ear including the lesion and adjacent skin was excised from the euthanized mouse and cleaned by shaking in Betadine followed by four rinses in sterile PBS. The lesion (up to ∼40 mm^3^) was excised from this piece and placed on a sterile surface. Two pieces of the lesion (up to ∼10 mm^3^ each) were used for implantation. Recipient NSG mice were anesthetized with isoflurane and injected intraperitoneally with meloxicam analgesic (Loxicam, 10 mg/kg; Norbrook Laboratories). An approximately 5 mm incision was made in the right flank skin and an opening was cleared subcutaneously using sterile forceps. A piece of ear lesion was inserted in the subcutaneous space, and the wound was closed with surgical glue (VetBond, 3M, #1469SB). Prominences caused by subcutaneous masses or surface papillomas were measured weekly starting 2 weeks post implantation using calipers (General Tools & Instruments, #147).

### Histology

#### Fixation, sectioning, and H&E staining

Tongues were either frozen in O.C.T compound (Tissue-Tek) or fixed in Surgipath Decalcifier I (to soften parts of the hyoid bone associated with the BoT; Leica Biosystems, Buffalo Grove, USA) overnight at 4°C twice, with fresh decalcifier for the second incubation. Images show paraffin sections unless “frozen” is specified. Anuses, ears, lungs, and lymph nodes were fixed in 4% paraformaldehyde in PBS for 24-48 hours at 4°C. Lungs were filled with PBS or PFA prior to immersion in PFA. Fixed tissues were then transferred to 70% ethanol and processed. Processed tissues were embedded in paraffin and sectioned at room temperature using a manual rotary microtome (5 um sections; Leica RM2235). Frozen tongues were sectioned using a cryostat (10 um sections; Leica CM1950). Tongues were cut in half and sectioned sagittally (3 sections per slide, 50 slides). Anal tracts were cut in half lengthwise prior to sectioning along the luminal axis (3 sections per slide, 20 slides). The infected area of ears was trimmed, cut in half, and embedded on edge for cross-sectioning (3 sections per slide, 50 slides). Lymph nodes were embedded without directionality together with lungs, which were sectioned coronally (1 section per slide, 50 slides). Every 10^th^ section was stained with hematoxylin and eosin (H&E) as described^24^.

#### Immune labeling

For CD4 and CD8 T cell labeling, frozen sections were fixed by immersion in cold methanol and labeled with anti-CD4 (clone RM4-5, eBioscience), anti-CD8 (clone 53-6.7, eBioscience) and fluorescent secondary antibodies as described^32^. KRT14 and L1 capsid protein co-labeling of paraffin-embedded sections, with tyramide signal amplification of L1 staining, was performed as described^22^ – except anti-K14 (905301, Biolegend) was used at 1:5000. KRT14 and phospho-S6 co-labeling was performed in the same manner, including tyramide signal amplification, with anti-pS6 replacing anti-L1 (1:4000; Cell Signaling CS-4858L). KRT14 and TRP63 co-labeling of paraffin-embedded sections was performed after antigen retrieval with Antigen Unmasking Solution (pH 9, Vector Laboratories), using anti-KRT14 as above and anti-TRP63 (1:100; MAB 4135, Millipore), followed by fluorescent secondary antibodies (1:1000 donkey anti-rabbit AlexaFluor 594, A21207, Molecular Probes; 1:1000 goat anti-mouse AlexaFluor 488, A11001, Molecular Probes); slides were coverslipped with Prolong Diamond mounting media (P36970, Fisher Scientific). Ki67 immunohistochemistry was performed as described (King, Ward-Shaw, et al., 2022). Images were taken using a Zeiss Imager.M2 microscope with the addition of an EXFO X-Cite Series 120Q fluorescence illuminator for immunofluorescence images.

### RNA/DNA *in situ* hybridization

MmuPV1 nucleic acid was detected using RNAscope [15] 2.5 HD Assay-BROWN (#322300, Advanced Cell Diagnostics) according to the manufacturer’s instructions. Paraffin sections (5 µm) were hybridized with probes specific for MmuPV1 E4 or E6/E7 or with a negative control probe (MusPV-E4 #473281; MusPV-E6-E7 #409771; negative control #310043; Advanced Cell Diagnostics). To distinguish viral transcript from viral DNA, an adjacent section on the same slide was incubated for 30 min prior to probe hybridization with 20 U DNase I (EN0521, Fisher Scientific) and remaining sections were incubated with buffer only. One section on each slide was incubated either with no probe (NANOpure water, Barnstead) or with negative control probe.

### T cell depletion

For *in vivo* depletion of T cells, 100 ug each of antibodies to CD4 (BioXCell, clone GK1.5) and CD8 (BioXCell, clone 2.43) or 100 ug of isotype control (FVB depletion experiment) or 200 ug isotype control (FVB-E5 depletion experiment; BioXCell, Rat IgG2b, κ) was injected intraperitoneally twice weekly throughout the study, starting 4 days before MmuPV1 infection. Flow cytometry to assess levels of CD4 and CD8 T cells was performed using blood samples collected retro-orbitally just prior to tissue collection, as described^32^.

### Endoscopy

Endoscopy was performed using a 1.9 mm 0° endoscope in a 2.5 mm operating sheath (Karl Storz, El Segundo, USA) while mice were anesthetized in Mickey’s Space Helmet, as described^22^.

### Pathology

Hematoxylin- and eosin-stained slides from all tissues were assessed by experienced pathologists (R.H., J.P.S., D.B., K.M.) blinded to treatment group and timepoint post treatment. Reports included the presence of dysplasia (including severity) and cancer. The presence of inflammation and koilocytes was also reported for the base of tongue, and the presence of inflammation was reported for one ear infection experiment.

### MmuPV1 sequencing

MmuPV1 viral DNA was isolated from an aliquot of virus prep as follows. For DNA release from virion, 20 µl of virus (Prep 3) was incubated with 20 µl of viral release buffer (0.025 M EDTA, 0.5% SDS, 2 mg/ml Proteinase K) at 55°C for 1 hour. DNA was then purified using Qiagen’s PCR Purification Kit. DNA was used as template for two polymerase chain reactions (PCRs). The first reaction amplified the entire viral genome using primers that sit head to tail (5’-cttctgcaggatcttagctttgtctgc-3’; 5’-cagtgactcgaatgctttcaccgagtcgtctcc-3’; Integrated DNA Technologies). The second reaction amplified the region to which the first set of primers annealed, to provide complete coverage of the viral genome (5’-tggaaatcggcaaaggctacactc-3’; 5’-agccccaaacacagctacgaccc-3’). Both PCR products were gel purified and isolated using Qiagen’s Gel Extraction Kit and sent to PlasmidSaurus for sequencing. Sequence was visualized and compared to published MmuPV1 sequence (Joh et al., 2010) using SnapGene software (GSL Biotech).

### Statistics

The significance of differences in disease severity between groups of mice was determined using Mstat software (Mstat version 6.5.1, McArdle Laboratory for Cancer Research; https://mcardle.wisc.edu/mstat/). Disease severity was weighted (e.g. no dysplasia=0; mild dysplasia=1; moderate-to-severe dysplasia=2; invasive squamous cell carcinoma=3), and the set of numbers representing the disease severity of all mice in an experimental group was entered as a variable. These sets were compared using a two-sided Wilcoxon Rank Sum Test. The significance of differences in cancer frequency was assessed using Fisher’s Exact test. The significance of differences in the severity of disease across time or dose was assessed by chi-square analysis and confirmed by ordinal regression analysis using the “polr” function within the “MASS” R package (Venables and Ripley, 2002).

## Acknowledgements

The authors acknowledge Jim Murray, Bill Sugden, Caroline Alexander, Magdalena Murray, Bill Dove, Norman Drinkwater, Jenny Gumperz, and the Lambert laboratory for thoughtful discussions and insights; Wei Wang for sharing immunological expertise; Thomas Pier and the UW Experimental Animal Pathology Laboratory for tissue processing; Katherine Fox and the UW Flow Cytometry Laboratory for help with sorting and analysis; Susan Thibeault for sharing endoscopy equipment; and Patricia Esser, UW Biomedical Research Model Services, and UW Research Animal Resources and Compliance for expert animal care involving infectious disease. This project was supported by an American Hair Research Society Mentorship Grant and a UW Academic Staff Professional Development Grant (AB); NIH NIDCD F31 DC018184 and NCI T32 CA090217 (REK); NIH grants P01 CA022443 and R35 CA210807 (PFL); Jackson Laboratory Cancer Center Support Grant P30 CA089713; Specialized Program of Research Excellence (SPORE) program, through the National Cancer Institute grant P50CA278595; and the University of Wisconsin Carbone Cancer Center Support Grant P30 CA014520. The content is solely the responsibility of the authors and does not necessarily represent the official views of sponsoring institutions.

## Author contributions

A.B. and P.F.L. designed experiments, with contributions from R.E.K. (endoscopy), D.L.L. (virus sequencing), D.B. (long-term cutaneous infection), and R.H. (analysis of grafts and metastases). A.B. performed animal experiments, virus preparation, in situ hybridization, immunohistochemistry, immunofluorescence, flow cytometry (with assistance from the UW Flow Cytometry Laboratory), microscopic imaging, and WRS and FE statistical tests. E.T.W-S embedded and sectioned tissues and performed H&E staining. D.L.L performed virus purification, amplification, and sequence analysis. R.E.K. performed endoscopy and assisted with virus preparation and tissue collection. M.A.N. performed Chi-square and ordinal regression statistical analyses. D.B., K.A.M., J.P.S., and R.H. performed histopathological analysis. A.B. and P.F.L. prepared the manuscript with contributions from all authors.

## Competing interests

Sales of Mickey’s Space Helmet are licensed through the Wisconsin Alumni Research Foundation. A.B. and P.F.L. are inventors of Mickey’s Space Helmet and receive a portion of the proceeds. All other authors have no competing interests.

## Extended Data

**Extended Data Table.** Viral in situ hybridization and immunofluorescence.

| Tissue | Strain | Timepoint | Treatment | Severity | E6/E7 DNA/RNA | E6/E7 RNA | E4 DNA/RNA | L1 |
| --- | --- | --- | --- | --- | --- | --- | --- | --- |
| Base of Tongue | FVB | 2 weeks | MmuPV1 | Mild (4); Mod (2); SCC (2) | NP; 1/1; 2/2 | NP; NP; 2/2 | 4/4; 2/2; 2/2 | NP; NP; 1/2 |
| Base of Tongue | FVB-E5 | 2 weeks | MmuPV1 | No Dysplasia (3); Mild (2); SCC (5) | NP; NP; 2/2 | NP; NP; 2/2 | 1/3; 2/2; 5/5 | NP; NP; 2/2 |
| Oral Tongue | FVB-E5 | 2 weeks | MmuPV1 | Moderate Dysplasia | 2/2 | 2/2 | 2/2 | NP |
| Anus | FVB-E5 | 2 weeks | MmuPV1 | SCC | 2/2 | 2/2 | 2/2 | 2/2 |
| Ear | FVB-E5 | 2 weeks | MmuPV1 | Moderate Dysplasia (2); SCC (2) | 2/2; 2/2 | 2/2; 2/2 | 2/2; 2/2 | 2/2; 1/2 |
| Base of Tongue | NSG | 2 weeks | MmuPV1 | Mild Dysplasia | NP | NP | 1/1 | NP |
| Base of Tongue | B6 | 1 week | MmuPV1 | No Dysplasia (1); Mild Dysplasia (3) | 0/1; 3/3 | NP | NP | NP |
| Base of Tongue | B6 | 2 weeks | MmuPV1 | No Dysplasia | 2/8 | NP | 2/8 | NP |
| **T-cell Depletion** |  |  |  |  |  |  |  |  |
| Base of Tongue | FVB-E5 | 2 weeks | MmuPV1 + control Abs | SCC | 1/2 | 1/2 | 0/1 | NP |
| Base of Tongue | FVB-E5 | 2 weeks | MmuPV1 + T-cell Abs | SCC | 1/2 | 0/2 | 0/1 | NP |
| Base of Tongue | FVB-E5 | 4 weeks | MmuPV1 + control Abs | SCC | 1/1 | 1/1 | NP | NP |
| Base of Tongue | FVB-E5 | 4 weeks | MmuPV1 + T-cell Abs | SCC | 2/3 | 2/3 | NP | NP |
| Base of Tongue | FVB-E5 | 4 weeks | MmuPV1 | SCC | 2/2 | 2/2 | NP | NP |
| Base of Tongue | FVB | 2 weeks | MmuPV1 + control Abs | SCC | 1/1 | 1/1 | NP | NP |
| Base of Tongue | FVB | 2 weeks | MmuPV1 + control Abs | No Dysplasia | 4/4 | 4/4 | NP | NP |
| Anus | FVB | 2 weeks | MmuPV1 + T-cell Abs | SCC | 1/1 | 1/1 | 1/1 | 1/1 |
| Anus | FVB | 4 weeks | MmuPV1 + control Abs | SCC | 0/1 | 0/1 | 0/1 | 0/1 |
| **Timecourse** |  |  |  |  |  |  |  |  |
| Base of Tongue | FVB-E5 | 4 weeks | MmuPV1 | SCC (3) | 3/3 | 3/3 | NP | NP |
| Base of Tongue | FVB-E5 | 8 weeks | MmuPV1 | Severe (1) | 1/1 | 1/1 | NP | NP |
| Ear | FVB | 2 weeks | MmuPV1 | Mild (1); Severe (1); SCC (4) | 1/1; 1/1; 4/4 | 1/1; 1/1; 4/4 | NP; 1/1; 3/3 | NP; 1/1; 3/3 |
| Ear | FVB | 4 weeks | MmuPV1 | No Dysplasia (2); Mild (1); Poss SCC (1) | 2/2; 1/1; 1/1 | 2/2; 1/1; 1/1 | NP | NP |
| Ear | FVB | 8 weeks | MmuPV1 | Severe | 4/4 | 4/4 | NP | NP |
| Ear | FVB-E5 | 2 weeks | MmuPV1 | Mild (1); Mod (2); Poss SCC (3); SCC (2) | 1/1; 2/2; 3/3; 2/2 | 1/1; 2/2; 3/3; 2/2 | NP; 2/2; NP; 2/2 | NP; 2/2; NP; 1/2 |
| Ear | FVB-E5 | 4 weeks | MmuPV1 | Poss SCC | 4/4 | 4/4 | NP | NP |
| Ear | FVB-E5 | 8 weeks | MmuPV1 | Severe (2); Poss SCC (2) | 2/2; 2/2 | 2/2; 2/2 | NP | NP |
| Ear | FVB-E5 | 24 weeks | MmuPV1 | Poss SCC (1); SCC (3) | 1/1; 3/3 | 1/1; 3/3 | NP | NP |

**Extended Data Figure 1.**
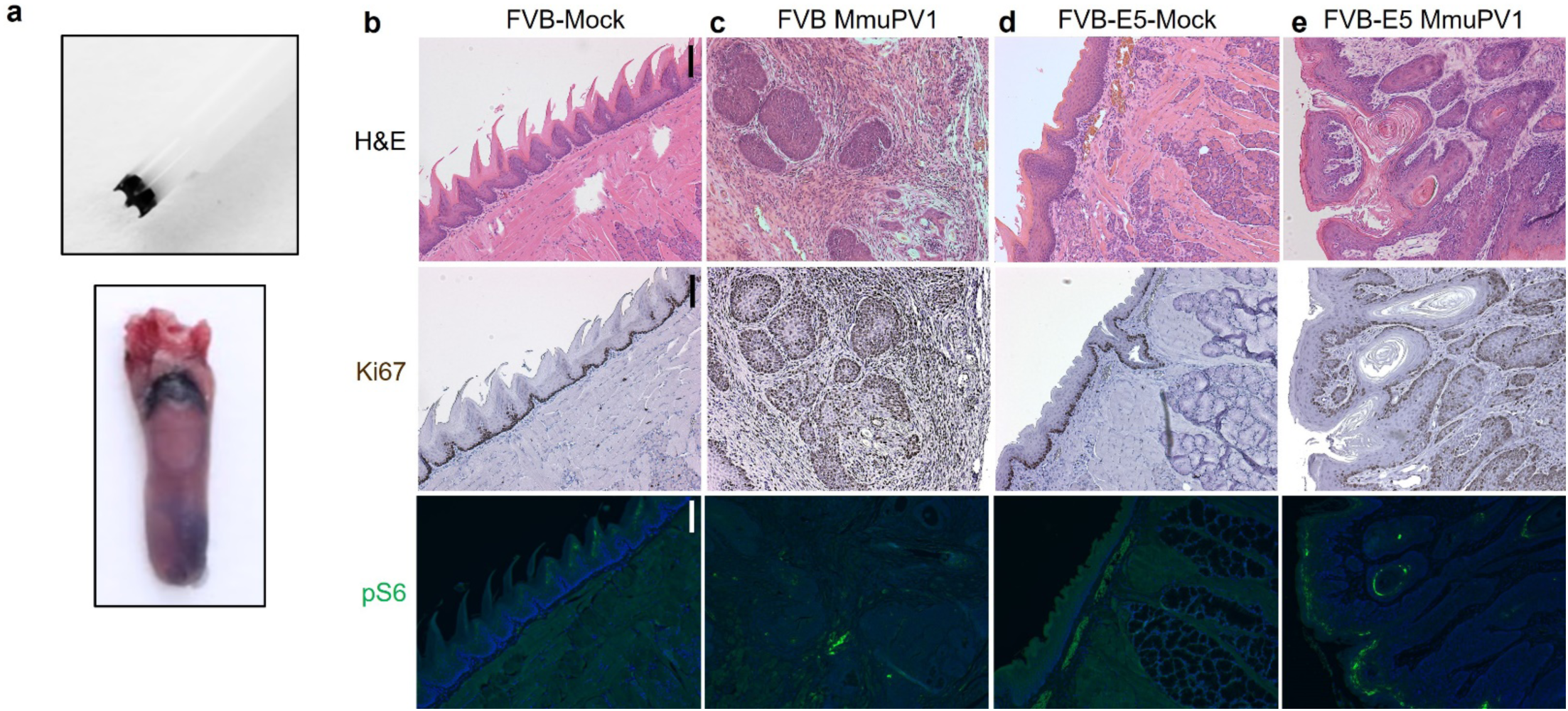
Cancers at the base of tongue. **a,** Top: A Greer Pick, shown holding approximately 1.5 ul 5% Evans Blue. Bottom: A mouse tongue that was mock-infected with 5% Evans Blue at the base of the tongue. **b-e,** Columns represent serial sections from a single mouse infected with ∼10^9^ VGE of MmuPV1 or mock-infected with PBS at the base of the tongue. Top row: stained with H&E; center row: labeled with antibody to Ki67 and DAB; bottom row: labeled with antibody to phospho-S6 and fluorescent secondary (green). Scale bar=100 um. **b,** mock-infected FVB, no dysplasia; **c,** infected FVB, invasive cancer; **d,** mock-infected FVB FVB-E5, no dysplasia; **e,** infected FVB-E5, invasive cancer.

**Extended Data Figure 2.**
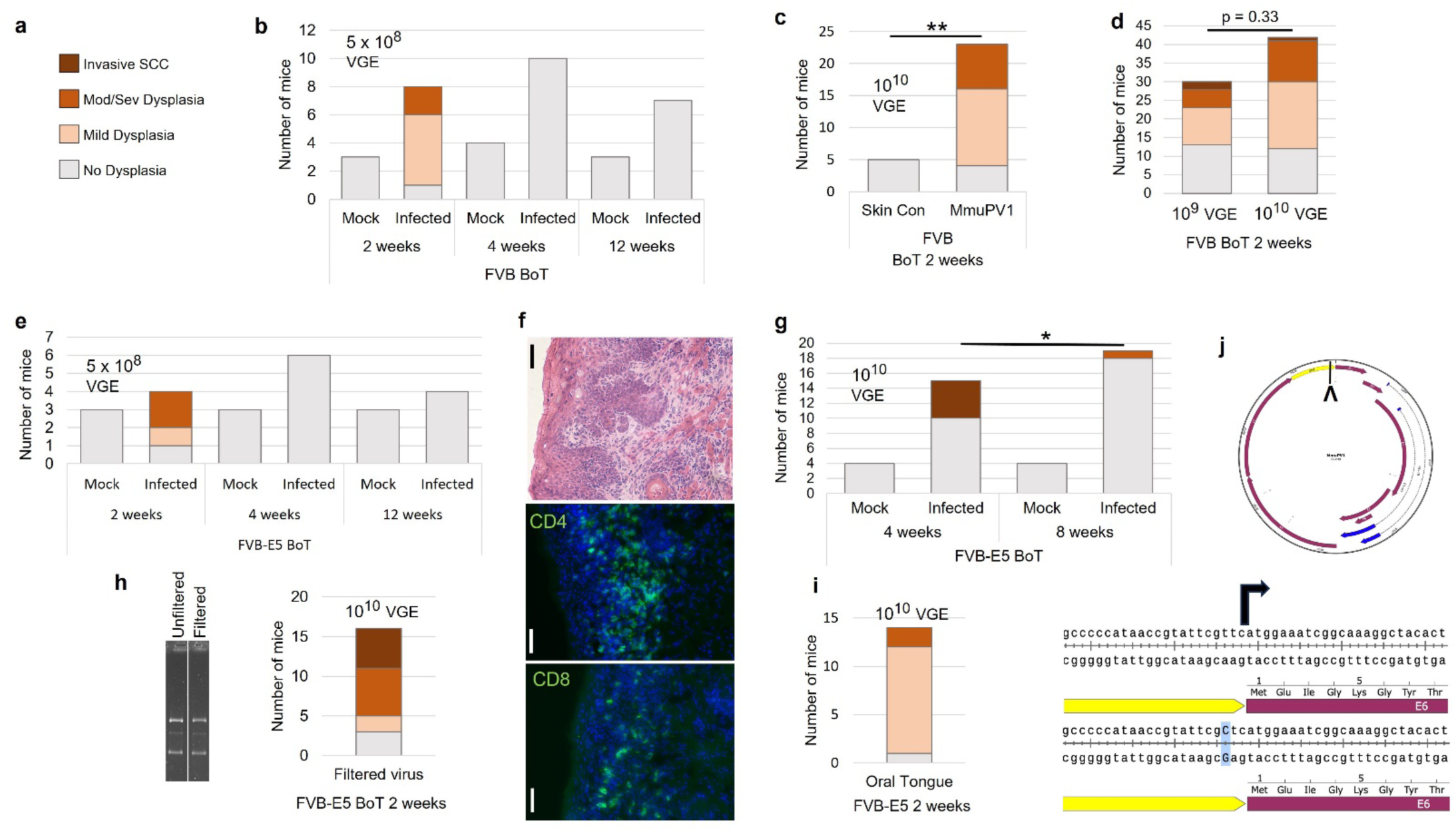
Cancers at base of tongue: additional doses and filtration of virus; timecourses; T cell infiltration; oral tongue infection; viral sequence. **a,** Legend for graphs in **b-e**, **g-i**. **b,** Lesion severity in FVB mice infected at base of tongue with ∼5 x 10^8^ VGE and collected at the indicated timepoints. **c,** Lesion severity in FVB mice infected with ∼10^10^ VGE (or mock infected with a corresponding control skin prep) at the base of tongue. WRS test of difference in disease severity: p<10^-2^. **d,** Comparison of lesion severity in FVB mice infected at the base of tongue with either ∼10^9^ or ∼10^10^ VGE MmuPV1 and collected 2 w.p.i. Results at each dose reflect combined results from multiple experiments, including isotype controls from T-cell depletion experiment and infections involving different virus preps. WRS test of difference in lesion severity: p=0.33. **e,** Lesion severity in FVB-E5 mice infected at base of tongue as in **b**. **f,** FVB-E5 invasive cancer at base of tongue infected with ∼10^9^ VGE of MmuPV1. Panels show serial frozen sections; scale bar=50 um. Top panel: H&E staining; center and bottom panel: labeled with antibodies to CD4 (center panel) or CD8 (bottom panel) and fluorescent secondary antibody (green) and counter-stained with Hoechst nuclear stain (blue). **g,** Lesion severity in FVB-E5 mice infected at the base of tongue with ∼10^10^ VGE and collected 4 or 8 w.p.i. FE test of difference in cancer frequency, 4 weeks vs. 8 weeks: * p=0.011. **h,** Left: agarose gel showing virus before and after passage through a 0.23 um filter; lanes were excised from one picture of a gel in which both samples were run. Right: Lesion severity in FVB-E5 mice infected at base of tongue with ∼10^10^ VGE MmuPV1 that had been filtered. **i,** Lesion severity in FVB-E5 mice infected in the oral/anterior tongue with ∼10^10^ VGE and collected 2 w.p.i. **j,** Top: Illustration of the MmuPV1 genome, with a caret indicating a sequence difference between virus prep 3 (see Materials and Methods) and the sequence of MmuPV1 submitted to NCBI by Joh *et al.* (2010). Bottom: The sequence reveals a single change in the 5’ untranslated region of the E6 gene: substitution of a C for a T in the “Kozak” sequence, where an A or G is optimal for maximal expression (Kozak, 1986).

**Extended Data Figure 3.**
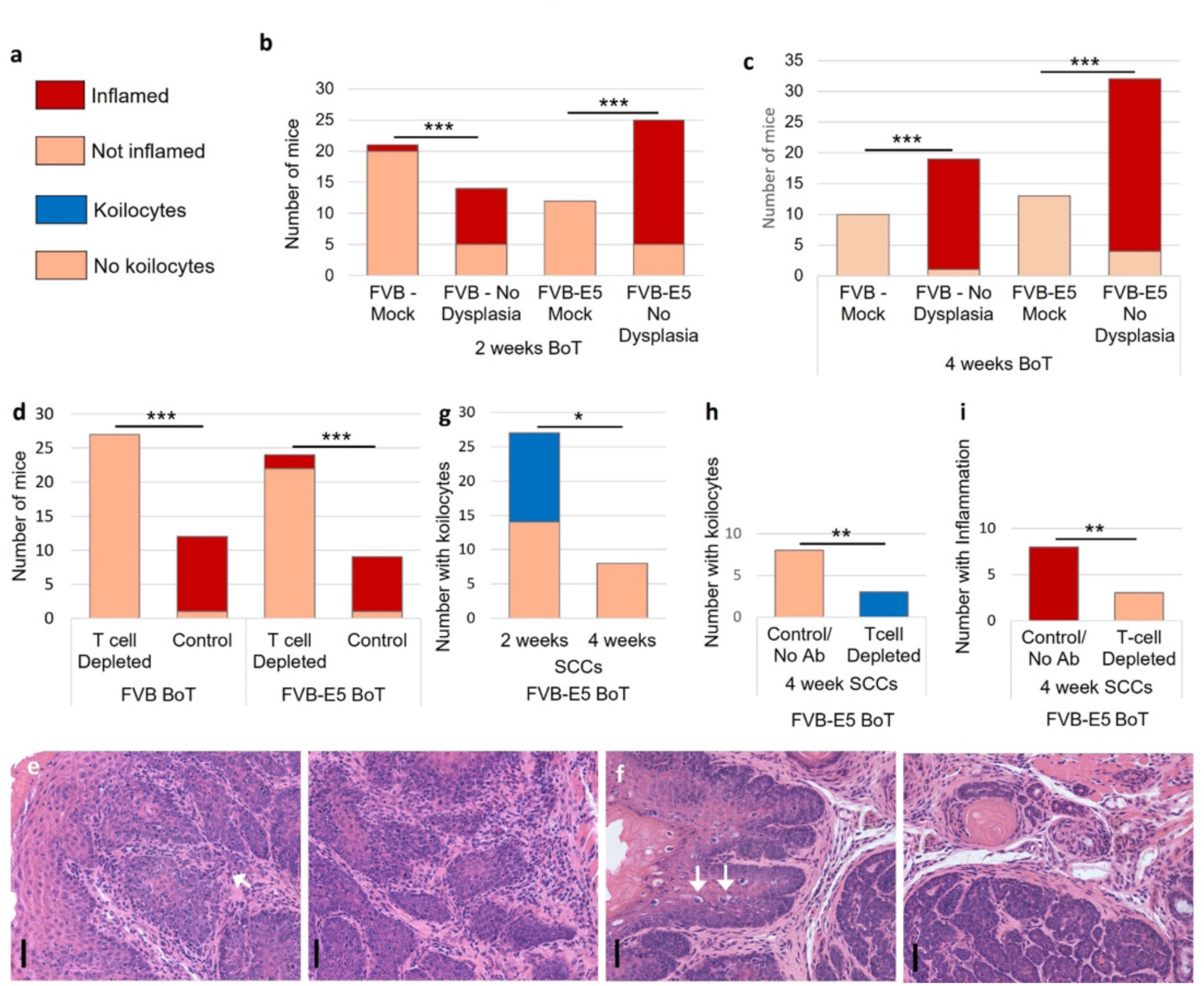
Inflammation and koilocytes at base of tongue. **a,** Legend for graphs. **b,c,** Inflammation in FVB and FVB-E5 mice either mock-infected or treated with MmuPV1 but displaying no dysplasia 2 weeks (**b**) or 4 weeks (**c**) post treatment at BoT (combined experiments, virus doses). **b,** FVB: *** p<10^-6^; FVB-E5 *** p<10^-7^. **c,** FVB: *** p<10^-3^; FVB-E5 *** p<10^-5^. **d,** Inflammation at base of tongue in dysplasia and cancer in FVB and FVB-E5 mice infected with ∼10^10^ VGE MmuPV1 and treated with either control or αCD4 and αCD8 antibodies. FVB: *** p<10^-8^; FVB-E5: *** p<10^-4^. **e,f,** High-magnification images of H&E-stained lesions 4 w.p.i., shown in 2**i**. Scale bar=50 um. **e,** Left panel: Image of surface epithelium and adjacent cancer tissue. No koilocytes, which are normally found in the upper epithelial layers, are visible. Inflammation is visible in both panels (arrow points to cluster of immune cells). **f,** Left panel: Image of surface epithelium with koilocytes (arrows point to 2 of several) and cancer. No inflammation is visible in this image or in adjacent area of cancer in right panel. **g,** Presence of koilocytes in FVB-E5 base of tongue SCCs 2 or 4 w.p.i. FE test of significance, * p=0.015. **h,i,** Presence of koilocytes or inflammation in FVB-E5 invasive SCCs 4 weeks after treatment with either control antibodies or no antibodies, or with αCD4 and αCD8 antibodies. FE test of significance. **h,** Koilocytes. ** p<10^-2^. **i,** Inflammation. ** p<10^-2^.

**Extended Data Figure 4.**
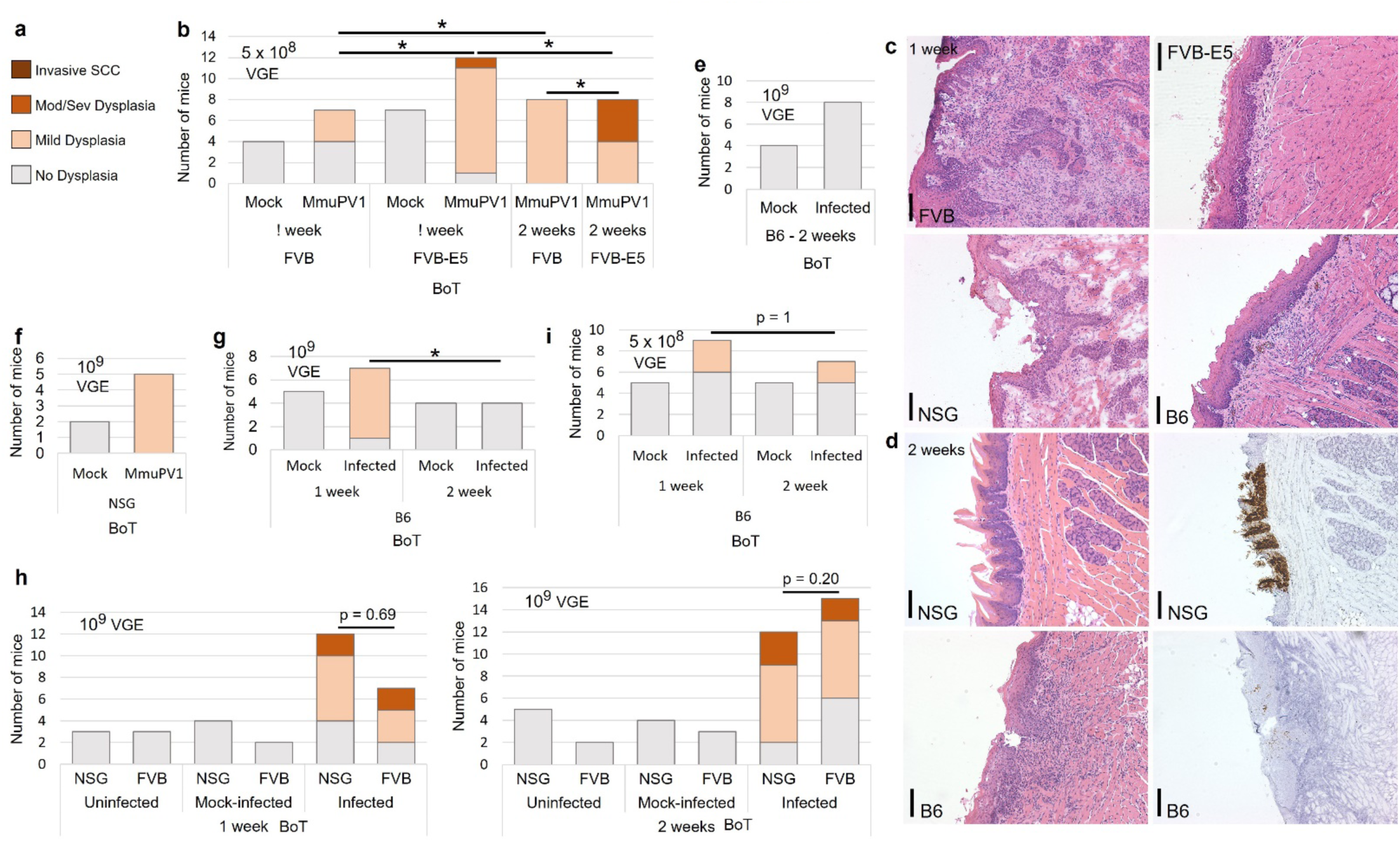
Base of tongue dysplasia develops within one week and persists to 2 weeks in a strain-dependent manner. **a,** Legend for bar graphs in **b**,**d-h**. **b,** Lesion severity in FVB and FVB-E5 mice infected with ∼5 x 10^8^ VGE or mock infected at the base of tongue with PBS, collected one or two weeks post infection. WRS test of difference in severity between infected mice: FVB vs E5 at 1 week, * p=0.023; FVB vs E5 at 2 weeks, * p=0.027; FVB 1 week vs 2 weeks, * p=0.016; E5 1 week vs weeks, * p=0.036. **c,d,** Scale bar=100 um. **c,** Base of tongue infected with MmuPV1 and collected one week post infection, stained with H&E. FVB, NSG (frozen sections): ∼10^9^ VGE, from experiment graphed in **h**; FVB-E5, ∼5×10^8^ VGE, from experiment in **b**; B6, ∼10^9^ VGE, from experiment in **g**. Panels: At least severe dysplasia, suspicious for early invasion, in FVB; mild dysplasia in FVB-E5; severe dysplasia in NSG; mild dysplasia in B6. **d,** Base of tongue infected with ∼10^9^ MmuPV1, collected two weeks after infection, stained with H&E (left panels) or RNAscope with MmuPV1 probe (NSG: E6/E7; B6: E4; right panels). Top panels: NSG, mild dysplasia; bottom panels: B6, no dysplasia. **e,f,** Lesion severity in mice collected 2 weeks after infection with ∼10^9^ VGE or mock infection at the base of tongue with PBS. **e,** B6. **f,** NSG. **g,** Lesion severity in B6 mice collected 1 or 2 weeks post infection with ∼10^9^ VGE FE test of lesion frequency, 1 week vs. 2 weeks: * p=0.015. **h,** Lesion severity in FVB and NSG mice infected with ∼10^9^ VGE one week (left panel) or two weeks (right panel) post infection. WRS test of difference in severity between infected mice: FVB vs NSG 1 week, p=0.69; FVB vs NSG 2 weeks, p=0.20; FVB 1 week vs. 2 weeks, p=0.48; NSG 1 week vs 2 weeks, p=0.40. **i,** Lesion severity in B6 mice infected with 5 x 10^8^ VGE 1 or 2 weeks post infection. FE test of difference in lesion frequency, 1 week vs. 2 weeks: p=1.

**Extended Data Figure 5.**
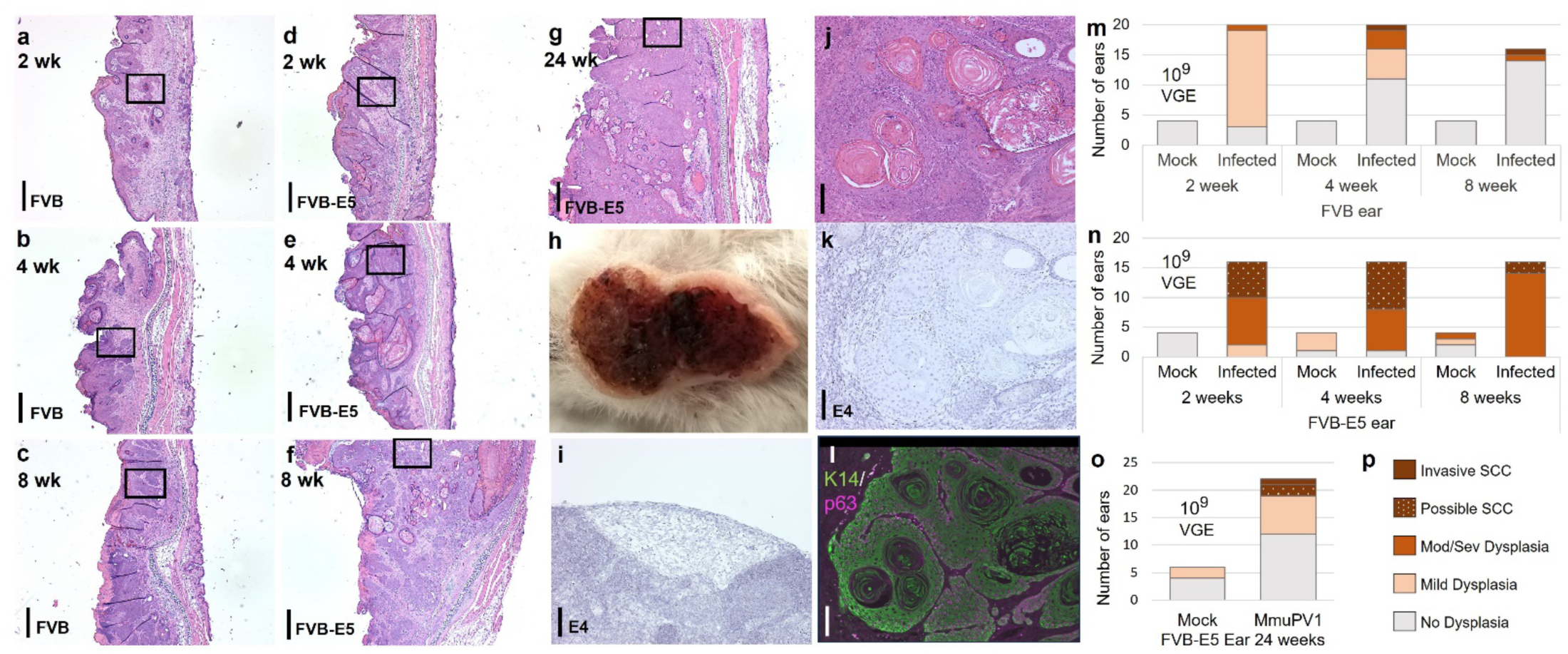
Ear lesions. **a-g,** H&E-stained cross-section of ears infected with ∼10^10^ VGE MmuPV1 and collected at the indicated timepoints. Scale bar=400 um. Area in boxes shown at higher magnification in Figure 4. **a-c,** FVB at 2, 4, and 8 weeks; **d-g,** FVB-E5 at 2, 4, 8, and 24 weeks. **h,** Left ear lesion in FVB-E5 mouse 38 w.p.i. with metastasis to lymph node. (Right ear shown in Fig. 4m.) **i,** Lymph node metastasis labeled using RNAscope with probe for MmuPV1 E4 DNA/RNA. Scale bar=50 um. **j-l,** Serial sections of invasive ear lesion that had no detectable MmuPV1 DNA/RNA (circled in Fig. 4r). Scale bar=50 um. **j,** H&E stain. **k,l,** Sections labeled using RNAscope with probes for MmuPV1 E4 DNA/RNA (**k**) or with antibodies to KRT14 and TRP63 (**l**). **m,n,** Severity of lesions at indicated time post infection of each ear with ∼10^9^ VGE or mock infection with control skin prep. **m,** FVB. **n,** FVB-E5. **o,** Severity of FVB-E5 ear lesions 24 weeks post infection with ∼10^9^ VGE MmuPV1. **p,** Legend for graphs in **m-o**.

**Extended Data Figure 6.**
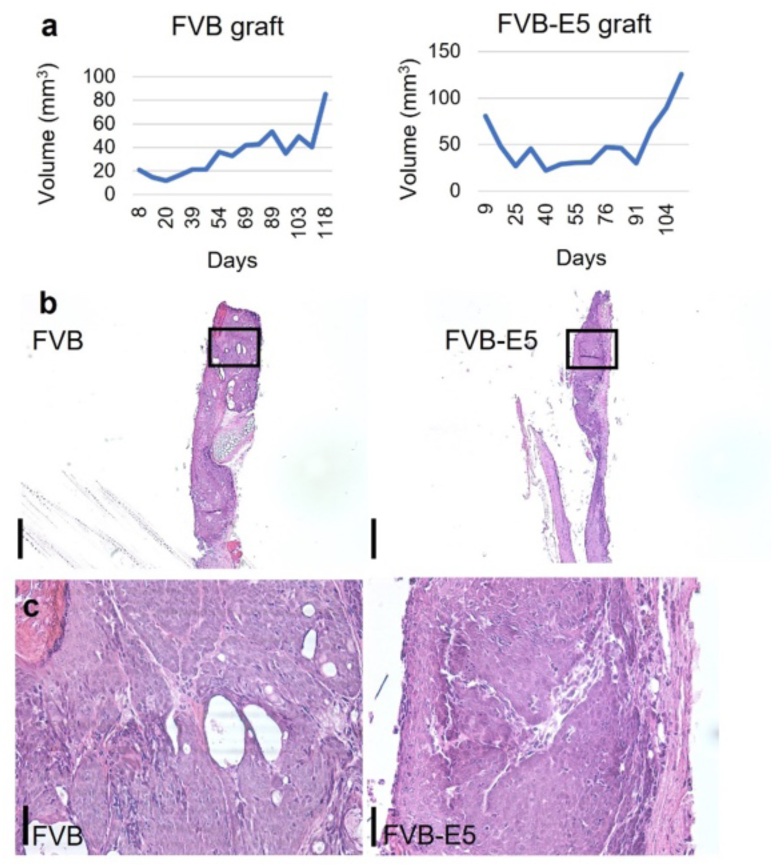
Ear lesions grow ectopically in an immune-deficient host. **a-c,** Ear lesion grafts. Ear lesions were collected two weeks post infection with ∼10^10^ VGE MmuPV1 and implanted subcutaneously in NSG mice. **a,** Grafts were measured weekly. Left panel: FVB graft; Right panel: FVB-E5 graft. **b,c,** Grafts were collected 17 weeks (left panel; FVB) or 16 weeks (right panel; FVB-E5) after transplantation of lesions; sections were stained with H&E. **b,** Scale bar=400 um; **c,** Sections of grafts shown boxed in **b; s**cale bar=50 um.

